# Constrained Instruments and their Application to Mendelian Randomization with Pleiotropy

**DOI:** 10.1101/227454

**Authors:** Lai Jiang, Karim Oualkacha, Vanessa Didelez, Antonio Ciampi, Pedro Rosa, Andrea L. Benedet, Sulantha Mathotaarachchi, J. Brent Richards, Celia M.T. Greenwood

## Abstract

In Mendelian randomization (MR), genetic variants are used to construct instrumental variables, which enable inference about the causal relationship between a phenotype of interest and a response or disease outcome. However, standard MR inference requires several assumptions, including the assumption that the genetic variants only influence the response through the phenotype of interest. Pleiotropy occurs when a genetic variant has an effect on more than one phenotype; therefore, a pleiotropic genetic variant may be an invalid instrumental variable. Hence, a naive method for constructing instrumental variables may lead to biased estimation of the causality between the phenotype and the response. Here, we present a set of intuitive methods (Constrained Instrumental Variable methods [*CIV*]) to construct valid instrumental variables and perform adjusted causal effect estimation when pleiotropy exists, focusing particularly on the situation where pleiotropic phenotypes have been measured. Our approach includes an automatic and valid selection of genetic variants when building the instrumental variables. We also provide details of the features of many existing methods, together with a comparison of their performance in a large series of simulations. *CIV* methods performed consistently better than many comparators across four different pleiotropic violations of the MR assumptions. We analyzed data from the Alzheimer’s Disease Neuroimaging Initiative (ADNI) Mueller et al. (2005) to disentangle causal relationships of several biomarkers with AD progression. The results showed that CIV methods can provide causal effect estimates, as well as selection of valid instruments while accounting for pleiotropy.

## 1 Introduction

### 1.1 Mendelian Randomization

Mendelian randomization is a method for estimating the causal effect of a modifiable exposure (**X**) on a disease (**Y**) by using measured genetic variation (**G**) as an instrument to eliminate bias from unmeasured confounding factors (**U**). The Mendelian inheritance patterns of genetic data, from parents to children, can be viewed as comparable to a randomized controlled trial. The reason is, if the choice of mate is not associated with genotype, the genotypes distribution (of the offspring) should be unrelated to any confounding factors. From a statistical perspective, Mendelian randomization is an application of instrumental variable methods using genetic information, as instruments (Smith and Ebrahim, 2004; Didelez and Sheehan, 2007; Lawlor et al., 2008; Wehby et al., 2008) for the exposure of interest **X**, as illustrated in Figure 1. In order to obtain valid results from instrumental variable analysis, several important assumptions about the relationships between the genotype instruments, **G** (usually single nucleotide polymorphisms (SNPs)), and the other variables must hold. When working with a structural equation modeling (SEM) set-up, the assumptions for Mendelian randomization are:

(A1) **G** is associated with the exposure **X** (i.e. 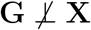, or **G** and **X** are not independent).
(A2) **G** and **Y** are independent conditional on exposure **X** and unmeasured confounding factors **U** (i.e. 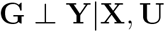).
(A3) **G** and confounders **U** must be independent (i.e. **G** ⊥ **U**).

**Figure 1:**
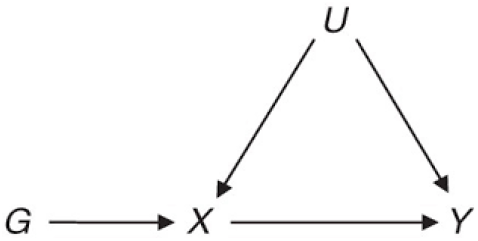
A directed acyclic graph (DAG) representing a situation where Mendelian randomization using genetic variants **G** as instruments can be useful for inferring whether a phenotype **X** is causally related to an outcome **Y**. **U** represents unmeasured confounding factors.

If linear models are assumed for the dependencies among the **G**, **X**, **Y**, then “independent” in the assumption can be relaxed to “uncorrelated”, and “associated” can be replaced with “correlated”. In the following we assume linear relationship for causal dependencies.

These assumptions may be violated in some contexts (Didelez and Sheehan, 2007; Lawlor et al., 2008). For example, linkage disequilibrium, which refers to the association of alleles at different loci in the population, may lead to violations of condition (A2). If the genetic variant of interest, Gı, is in linkage disequilibrium with another genetic variant, G_2_, which has a direct or indirect influence on the disease **Y**, then (A2) is not satisfied for *G*_1_. It is often believed that genotypes will not be associated with the socioeconomic and behavioral characteristics that confound **X** and **Y**, however, careful assessment of possible violations of (A3) is still necessary in some situations (Lawlor et al., 2008). Often most the problematic assumption is (A2), since genetic variants often have effects on cell functioning that are not well understood and could plausibly act through many mechanisms.

### 1.2 Challenges Arising from Pleiotropy

Pleiotropy-when more than one phenotype is influenced by the same group of genotypes-may violate assumption (A2) if all these phenotypes are themselves on the causal pathway for the response **Y**. Two kinds of pleiotropy can be defined (Solovieff et al., 2013): biological pleiotropy (Stearns, 2010; Wagner and Zhang, 2011) refers to associations involving multiple phenotypes sharing the common genetic pathways. For example, a gene variant in *PTPN22* is known to be associated with immune related disorders such as Type 1 diabetes (Todd et al., 2007) and Crohn’s disease (Barrett et al., 2008). This variant has been shown to interfere with the function of various T cells (Rieck et al., 2007) and affect the removal of autoreactive B cells (Menard et al., 2011). Hence, the impact of T cell levels on the risk of diseases will be confounded by the alternative causal pathway through reactive B cells, and vice versa. In contrast, mediated pleiotropy refers to direct causal impacts between phenotypes, such that the genotypes of interest have direct/indirect causal impact on both phenotypes. For example Thorgeirsson et al. (2008) reported a common variant in the nicotinic acetylcholine receptor gene cluster that affects both nicotine dependence (ND) and the smoking quantity (SQ). Both of these phenotypes (ND and SQ) are associated with the risk of lung cancer (Hung et al., 2008; Lamin et al., 2014) and yet smoking quantity can affect nicotine dependence. On the other hand, it is also believed that the most important factor for smoking persistence is nicotine dependence. The relationship between ND and SQ (Figure 2) depends on the duration of smoking and magnitude of craving (Donny et al., 2008). Therefore, in order to estimate the magnitude of the effect of nicotine dependence alone on lung cancer, one would need to appropriately account for the effect of smoking quantity. Methods to clarify such complex relationship are essential.

**Figure 2:**
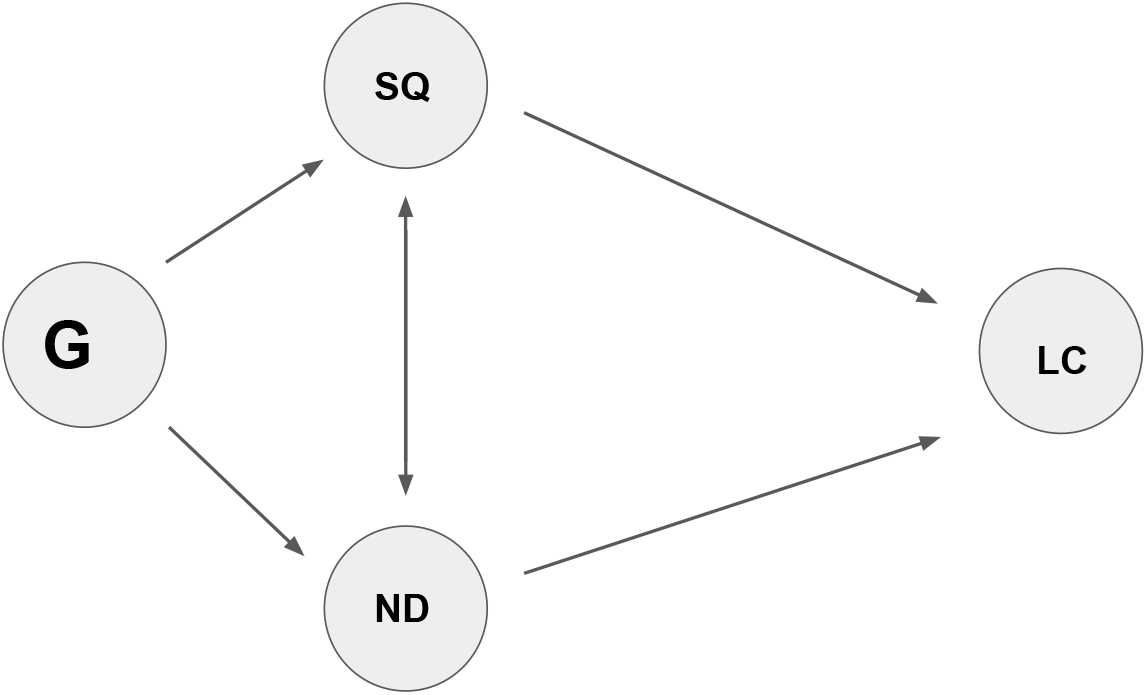
Diagram representing pleiotropy in smoking studies. **G**: genotypes. ND: nicotine dependence. SQ: smoking quantity. LC: lung cancer (risk). A bidirectional arrow between SQ and ND reflects multiple dependencies. For simplicity we omit possible confounding factors here.

The motivation for this work is derived from two straightforward questions:

(B1) In many applications, researchers may have access to large data collections including many phenotypes. Although some are of primary interest for their causal effects, others could be considered of secondary interest – yet may be influenced by some of the same genetic variants. What is the best way to use data from these additional phenotypes and perform inference on causal effects?
(B2) Does the solution for (B1) select genetic variants that *only* affect the phenotype of primary interest?

Many different statistical methods have been proposed to address challenges arising from pleiotropy. In section 2 we introduce notation and review several of the popular methods. Then in section 3, we propose a novel instrumental variable approach for Mendelian randomization in the presence of potential pleiotropic phenotypes. First, we describe our instruments, which are based on a weighted combination of the original genetic information and maximize the association between **G** and **X** under a set of constraints. We then show that approximate sparse solutions for these weights can be obtained, and furthermore that we can select approximately valid instruments as a result. In section 4, we compare all these methods against each other in simulated data, and in section 5 we analyze an Alzheimer’s disease dataset ADNI to demonstrate the performance of our *CIV* method as well as other instrumental variable methods.

## 2 Context of the Problem and Review of Existing Methods

For each individual *i* ∈ {1, …, *n*}, let *Y_i_* be the response of interest and let 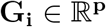 represents the genotypes that have been collected for *i*th observation, where *p* is the number of all genotypes available. Let 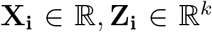 be the phenotypes that have been measured for this individual; **X_i_** is the phenotype of interest and **Z_i_** are phenotypes that may be affected by elements of **G_i_**. We denote 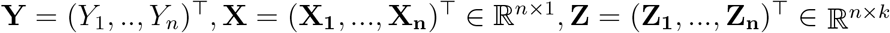 and 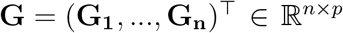 to be *n*-dimensional vectors while each individual is assumed to be observed in an i.i.d. fashion. We assume linear structural equation models:

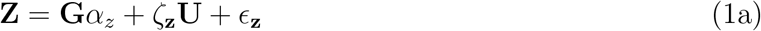

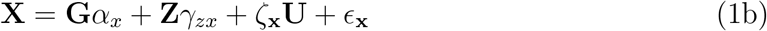

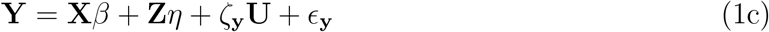

Or

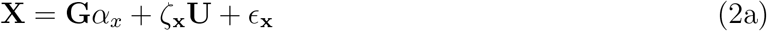

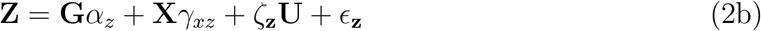

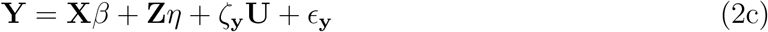

where *α_x_* and *α_z_* are the association parameters between genotypes **G** and phenotypes **X**, **Z**. *β* is the causal effect parameter of interest, and *η* is the pleiotropic causal effect of **Z** on **Y**. ζ_x_, ζ_z_, ζ_y_ represent the impact (coefficient) of unmeasured confounding factors **U** on **X**, **Z** and **Y** respectively. We use *γ_zx_* and *γ_xz_* to denote the direct causal impact of **Z** on **X** and **X** on **Z** respectively. Note that at least one of *γ_zx_* and *γ_xz_* should be 0. Let e_x_,e_z_,e_y_ be independent errors for **X**, **Z** and **Y** respectively.

Figure 3 lays out the general pleiotropic structure of this model. We assume that genotypes **G**, phenotypes of interest **X**, the response **Y** and potential pleiotropic phenotypes **Z** have all been measured for each individual in the study. The parameter of interest is assumed to be β, the causal effect of **X** on **Y**. Unobserved confounders are indicated by **U**. The relationships between phenotypes of interest **X**, pleiotropic phenotypes **Z**, and outcomes **Y** may vary from one situation to another, i.e. not all of the edges or arrows in Figure 3 need to be present in any particular example. However, if **X** has a direct causal relationship with **Y**, then *β* must be nonzero.

**Figure 3:**
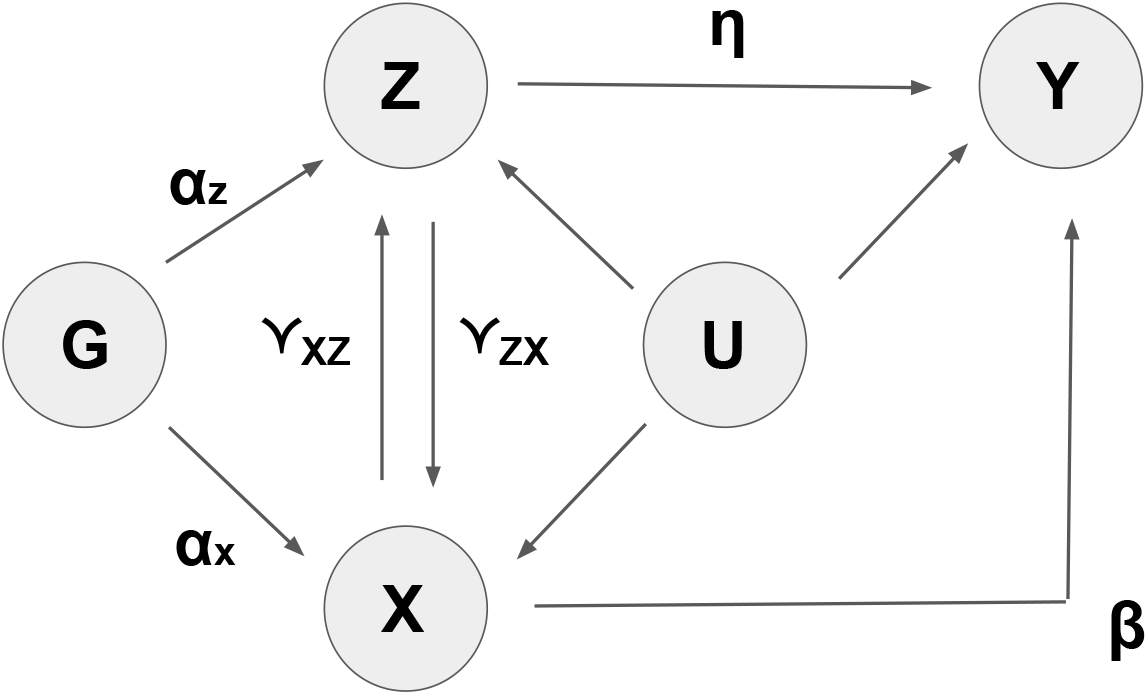
General diagram representing pleiotropic influences in Mendelian randomization studies. **G**: genotypes. **X**: phenotype of interest. **Z**: pleiotropic phenotypes. **Y**: response of interest, q**⅛,**q**⅛** are the genetic association parameters between **X** ~ **G** and **Z** ~ **G**. *β* is the causal effect of interest (**X** on **Y**). *η*: the pleiotropic pathways of **Z** on **Y**. *γ_xz_* and *γ_zx_* represent the possible causal effects of **X** on **Z** and **Z** on **X** respectively.

Three different types of relationship between **X** and **Z** can be assumed:

i. **X** and **Z** are conditionally independent given **G** and **U** (*γ_zx_* = *γ_xz_* = 0).
ii. direct causal impact of **X** on **Z** (*γ_xz_* ≠ 0 and *γ_zx_* = 0).
iii. direct causal impact of **Z** on **X** (*γ_zx_* ≠ 0 and *γ_xz_* = 0).

It is worth noting that only (i) and (iii) can be referred to as pleiotropy (by definition) when *α_z_* = 0. For (ii) even valid instruments (with *α_z_* ≠ 0) are still associated with **Z** due to the path **G** → **X** → **Z** and the total causal effect for **X** would be *β* + *γ_xz_η*.

The term “endogenous” variable describes the factors that are explained by the genotype-phenotype relationships and have impact on response **Y**. Common endogenous variables include health-related behaviors and risk-related phenotypes. **X** and **Z** in Figure 3 are both endogenous since they are both determined by genotypes and have impact on response, albeit with different functions. Variables such as age and sex that are not associated with the genotype-phenotype causal pathways of interest are termed “exogenous” variables; normally it is possible to adjust for these variables in a straightforward way in data analysis.

### 2.1 One Sample and Two Sample Mendelian Randomization

MR can be conducted with one sample of subjects, where individual level data (**G**, **X**, **Y**) or summary statistics (**G-X** associations and **G-Y** associations) are used to infer the causal effect **X** → **Y**, all in the same data set. An alternative strategy is to use two-sample MR methods with summary statistics, when no joint data is available. The gene-exposure (**G-X**) and gene-outcome (**G-Y**) associations are taken from different data sources for two-sample MR with summary statistics. Some MR methods for individual level data can also be separated into two steps and used with two-sample data. However, not all MR methods are straightforwardly adapted for two-sample set-ups.

There are three main reasons for using two-sample MR: the first is the separation of **G-X** and **G-Y** associations in two datasets. In this case only two-sample MR is applicable. Moreover, even if the estimated genetic instrument’s effect on **X** is biased in the first data set, this bias should not affect the causal effect estimation obtained in the second data set (Lawlor, 2016). The second reason is to alleviate weak instrument bias, in which instruments are weakly associated with phenotype of interest. As a result, in one-sample analysis, the (unobserved) confounders may explain more variation in the phenotype than the instruments, and the estimates will be biased towards the observational confounded association 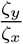 (Burgess et al., 2011). In two sample MR analysis the over-fitting of **X** is avoided, and then the weak instrument bias is towards the null (Davey Smith and Hemani, 2014). The third reason for using two-sample MR is the possibility to improve the statistical power of causal effect estimates and reduce **G** → **X** bias by using large data collections with many cases of samples.

### 2.2 Approaches based on 2SLS

If possible, a simple solution to address pleiotropy is to select for analysis only the valid genotypes, i.e. those that influence the phenotype of interest **X** and not **Z**, and use them as instruments with instrumental variable methods such as such as two-stage least squares (2SLS) regression. 2SLS method is a popular technique that is used in the analysis of structural equations. Given valid instruments **G**, the following two steps define a 2SLS model:

1. In the first stage, we obtain a new variable **X**\* as fitted value from ordinary least square regression **X** ~ **G**, where **G** are the selected instruments.
2. In the second stage, we substitute **X** with **X**\* and obtain ordinary least square estimators of *β* from the regression **Y** ~ **X**\*.

However, this selection of valid instruments may not always be possible without in-depth knowledge of the disease under study. For example, Timpson et al. (2005) uses common CRP (C-reactive protein) gene haplotypes, based on 3 SNPs, as instruments to infer the causal effect of the CRP protein (i.e. the phenotype of interest **X**) on multiple metabolic syndrome phenotypes including body-mass index (BMI) and high density lipoprotein (HDL) levels (i.e. **Y**. However, Martínez-Calleja et al. (2012) showed that one of these SNPs (rs1130864) is directly related to BMI, and it is also known that there exists a negative association between higher levels of BMI and HDL (Shamai et al., 2011). Hence, these 3 SNPs may not all be valid instruments for causal effect estimation of CRP protein levels on HDL. In many situations, our understanding of SNP effects is not complete enough to select valid instruments based on knowledge.

Another simple solution is to replace **G** with residuals after regressing each of these genetic variants on the pleiotropic phenotypes, **Z**. Specifically, one can replace **G** with **G**\* = (**I** − **P_z_**)**G** where 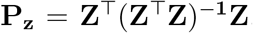, which regresses out the exogenous effect of **Z**. Then two-stage least squares (2SLS) or other instrumental methods can be applied to the new **G**\*, **X**, **Y**. We refer to this method as “2SLS_adj”. In general, this is not an appropriate solution and it will lead to biased estimation of *β* since **Z** is not an exogenous variable.

A related solution based on the underlying linear structural equation model (Equation 1) turns to multiple linear regression of **Y** on 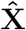 and **Ẑ** jointly in a two-stage least squares (2SLS) model, where 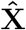 and **Ẑ** are the predicted phenotypes using **G** as the instruments. We refer to this method as *“2SLS-mul*” since it essentially implements multiple 2SLS regression. The *2SLS-mul* method uses **G** to account for the endogeneity of **Z** without controlling for it explicitly. However, using this approach, the resulting estimator of *β* will be unstable if 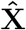 and **Ẑ** are highly correlated (Farrar and Glauber, 1967; Graham, 2003; Grewal et al., 2004).

By nature *2SLS-ãdj* and *2SLS-mul* methods require individual level data in a single sample. In the two-sample context, when (**G, X**) and (**G, Y**) are observed separately, the estimators are still available for 2SLS under certain conditions and assumptions (Angrist and Krueger, 1992; Dee and Evans, 2003). In that case, the corresponding estimators are called “two-sample instrumental variable” (TSIV) estimators in empirical studies (Borjas, 2004; Dee and Evans, 2003). In this paper, we only refer to *2SLS* methods as the estimators for individual level data.

### 2.3 Distribution of Causal Effects Across Multiple Instruments

When there are multiple instruments but not all are valid, causal conclusions can still be drawn by examining the distribution of the estimated causal effects across the instruments. If a group of genotype instruments, such as SNPs, are independent-i.e. located at different sites in the genome-and they all lead to similar estimates of the causal effect of **X** on **Y** (i.e. *β*), then this pattern provides strong evidence of a causal relationship between **X** and **Y**. This phenomenon is nicely illustrated with a funnel plot, where the precision of 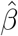 (defined as the reciprocal of its variance) for each SNP is plotted against its estimate, 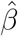. Asymmetry in a funnel plot might indicate an unbalanced pleiotropy, where variants subject to pleiotropy tend to bias the causal estimate in the same direction. There are tests for symmetry such as Beggs rank correlation test, although they have low statistical power (Begg and Mazumdar, 1994).

Mendelian randomization with *Egger* regression *(MR-Egger*) provides a more specific way to assess whether pleiotropy is present and to obtain an “unbiased” estimate. *Egger* regression is defined as the linear regression of a normalized parameter estimate against its precision (reciprocal of the corresponding standard error) in meta-analysis. *Egger*’s test assesses small study bias by testing the hypothesis that intercept of the *Egger* regression is zero. (Bowden et al., 2015) suggested that bias due to pleiotropy can be considered as analogous to small sample bias (Egger et al., 1997), and therefore that meta-analysis methods could be of use in Mendelian randomization settings. Mendelian randomization with *Egger* regression (MR-Egger) uses the slope coefficient from *Egger* regression as a consistent causal effect estimator of *β*. A key assumption (the InSIDE assumption) here is that the associations of the genetic variants with the phenotypes of interest are independent of the direct effects of the genetic variants on the response – i.e., *α_x_* is independent of *α_z_* for any SNPs in **G** (Figure 3). The InSIDE assumption still holds even if some instruments are invalid (*α_z_* ≠ 0), but it will be violated if genotypes **G** influence pleiotropic phenotypes (or confounders) **Z** that have impact on both **X** and **Y** – i.e., *α_x_* = 0,α_z_ ≠ 0,*γ_zx_* ≠ 0. MR-Egger regression provides no analytical estimate of the standard error of the *β*, although a confidence interval can be obtained using bootstrap methods. It is also worth noting that MR-Egger can be extended to two-sample summary data MR analysis. However, for individual level data analysis, MR-Egger is restricted to a single sample because it considers risk factors one at a time. If two distinct samples of **G**, **X**, **Y** are available, MR-Egger can be applied separately to both datasets and evaluates the difference between the two samples.

### 2.4 Summaries of Multiple Instruments

The genetic information across a set of genotypes can be summarized into a single genetic risk score by calculating a weighted sum, across the set, of the number of risk alleles carried by each person. The *Allele Score* method (Burgess and Thompson, 2013) is one example of this approach. Unweighted scores are simply the total number of risk-increasing alleles carried by an individual; weighted scores will allow the contribution of each risk-increasing allele to be proportional to the estimated additional risk. The weights are ideally derived from external information, such as previously published genetic associations, to reflect the corresponding genetic effect sizes. If no such external information exists, then cross-validation methods can be used to obtain bias-reduced weights for score construction, which then improves the causal effect estimation.

Several other weighted score methods have also been proposed. The Jackknife instrumental variable estimation method *(JIVE*) (Angrist et al., 1995) estimates 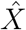 of each individual in the first stage regression *X* ~ *G* without using the corresponding data points. That is, the ith row in the jackknife matrix 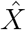 is estimated without using ith observation. As a result, that *JIVE* estimator is restricted to a single sample.

When the associations between instruments and the endogenous explanatory variable **X** are weak, the naive 2SLS estimator for *β* may be biased. All weighted score methods (including allele scores and *JIVE*), are expected to be able to reduce weak instrument bias in comparison to multiple instrument methods as described in section 2.3. In fact, weighted scores can be strong instruments even when the individual genetic variants are all weak instruments, and hence bias due to weak instruments is expected to be reduced. However, weighted scores usually also reduce the power (sensitivity) of MR studies (Pierce et al., 2010; Palmer et al., 2012) and require all instruments to be valid.

One limitation of this set of methods is that the choice of weights has a considerable impact on the bias of estimates. One popular and usually stable approach is to obtain weights from an external source with a large sample size, although this could introduce some bias due to cohort differences. If relevant external weights are not available, then we have to use “interval” weights derived from the data under analysis. However, there is no analytical standard error estimation of the resulting causal effect when internal weights are used.

In the context of two-sample analysis, the *Allele score* weights obtained from one sample can be used to infer the causal effect on a second distinct sample, given that **G**, **X**, **Y** are observed in both samples. The weights can be generated by (1) averaging the cross-validated weights from the training data; or (2) sampling from a normal distribution around the training weight with a chosen standard deviation (e.g. 0.01). The latter approximates the uncertainty in the estimation of weights from external data (Burgess and Thompson, 2013). In our two-sample analysis we refer to these two methods as *Allele* and *Allelesim* respectively.

### 2.5 Reducing Correlations between **G** and the Environment

A more sophisticated way to reduce bias due to pleiotropy is to set up a stringent condition for the estimate of *β* that reduces the correlation between **G** and any other factors that influence **Y** without going through **X**. Suppose that

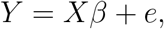

where *e* includes the effects of **U**, **Z** and *ϵ_y_*. Hence, the goal here is to reduce *Cor*(**G**, *e*), which is induced through the effects of **U** and **Z**. It must be noted that this only makes sense when there is no causal impact of **X** on **Z**. The Limited Information Maximum Likelihood (*LIML*, (Hayashi, 2000)) finds a conservative estimator of *β* by minimizing

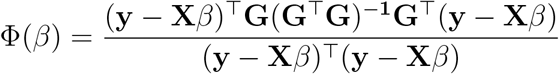

instead of minimizing 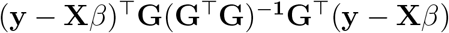 in the 2SLS. The denominator in the equation of Φ(*β*) is the variance of regression errors. *LIML* will be less biased than 2SLS when regressors **X** and regression error **Y** − **X***β* are not independent or when there are many weak instruments (Hahn and Inoue, 2002). However, the variance of *LIML* will increase when instruments are weak and the sample size is small, compared to 2SLS (Blomquist et al., 1999).

The idea behind *LIML* was generalized in the Continuously Updating Estimator approach (*CUE*, (Hansen et al., 1996)). This approach seeks “unbiased” estimator for 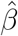 that satisfies the empirical analogue of the following (moment) conditions:

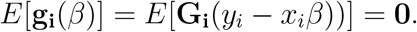

This is equivalent to minimizing

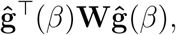

where 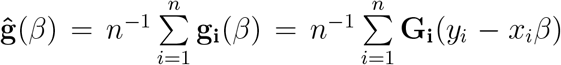 is the sample analog of the population moment conditions 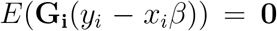 in the generalized method of moments (GMM) framework. Here, **G_i_** is the *i*th observation of instruments **G** and **W** is a weighting matrix. *CUE* defines a weighting matrix **W**(*β*) as a function 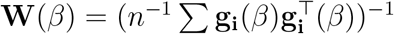 to give different weights to each moment condition.

All MR estimators based on the generalized method of moments (framework) are restricted to a single sample, since they all rely on the asymptotic properties of generalized methods of moments. Thus, given two distinct samples, *LIML* and *CUE* can only be applied separately to obtain one causal effect estimate for each sample.

Davies et al. (2015) demonstrated that *CUE* is a better choice than *LIML* and 2SLS when there are many weak instruments. In fact Hansen et al. (2012) and Newey and Windmeijer (2005) showed that generalized methods of moments estimators, including *LIML* and *CUE*, are more resistant to bias in this situation. Furthermore, Bound et al. (1995) showed that the bias of the 2SLS estimator can be approximated by:

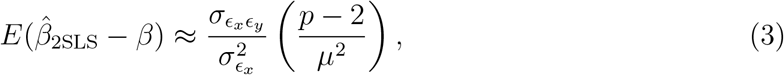

where 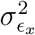 is the variance of the regression error *ϵ_x_* in the first stage, and *σ_ϵ_x_ϵ_y__* is the covariance of the second stage regression error *ϵ_y_* with *ϵ_x_*. The quantity 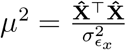 denotes the amount of the variation in **X** that is jointly explained by the instruments, where 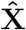 is the fitted value of **X** using **G**. Hence, the bias of the 2SLS estimator is proportional to the number of instruments (*p*) and inversely proportional to the variation in **X** that is explained by **G**. That is, the bias will increase if we add more instruments that do not explain the variation of **X**.

### 2.6 Valid Instrument Selection Methods

The correlation between **G** and 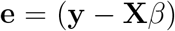 may arise from several kinds of violations of assumptions (A2) and (A3). The previously-described methods in sections 2.3, 2.4 and 2.5 do not use the information contained in **Z**. We have observed that when **X** and **Z** are highly correlated, both *LIML* and *CUE* fail to provide consistent estimates of *β*. Therefore, consistent solutions in the presence of pleiotropy have been proposed to embed valid instrument selection within an instrumental variable method. The motivation here is to incorporate both direct and indirect causal effects from **G** to **Y** using the following model:

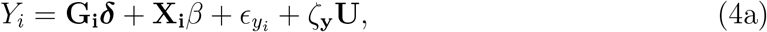

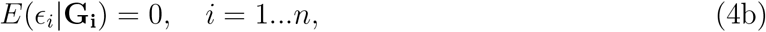

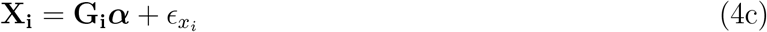

where ***δ*** represents the direct effects of the instruments **G** on outcome **Y**. Indirect effects of **G** on **Y** are captured through **X**, and *β* represents the causal effect parameter of interest. ***α*** is the association parameter between **G** and **X**. In recent work by Kang et al. (2016), the authors proposed the *some invalid some valid IV estimator* (sisVIVE) to minimize

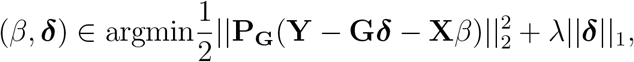

where 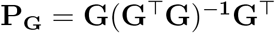. The first term essentially replaces **X** in 2SLS with (**X, G**), and thereby the direct causal effect of **G** on **Y** (via **Z**) is taken into consideration. The second term 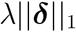 enforces an *L*_1_ sparse selection of invalid instruments using a LASSO prior. sisVIVE is robust to certain invalid instruments and their direct causal effect on **Y** (without going through **X**). However, this method’s ability to select valid instruments is limited by the assumption that at least 50% of all instruments must be valid.

sisVIVE treats pleiotropic phenotypes as general sources of the indirect causal effect **G** on **Y**, and does not use the information on **Z** explicitly. Given variables **Z**, Kang et al. (2016) suggests either adjusting for them or using them as exogenous variables. We refer to these two methods as *“sisVIVE*_adj” and *“sisVIVE*_exo” and implement them using R package sisVIVE.

If two distinct samples are available, sisVIVE can be extended to a two-sample estimator. First we can apply sisVIVE on a single sample to obtain estimation of ***δ*** and corresponding valid instrument selections. Then this estimation of ***δ*** can be carried over to the MR analysis of a second sample. The corresponding adjustment for pleiotropic phenotypes **Z** can also be incorporated within this two-sample estimator based on sisVIVE.

In summary, a high-level comparison of all the methods described above is provided in Table 1, contrasting key features. It is also worth noting that the descriptions above are largely restricted to the case of a univariate variable **X**, although most methods can be easily extended to multivariate **X** (see Discussion). Among all methods new approaches that can incorporate external information from a different sample (in two-sample setup), or construct valid instrumental variables to address pleiotropy, are attracting increased attention.

**Table 1:**
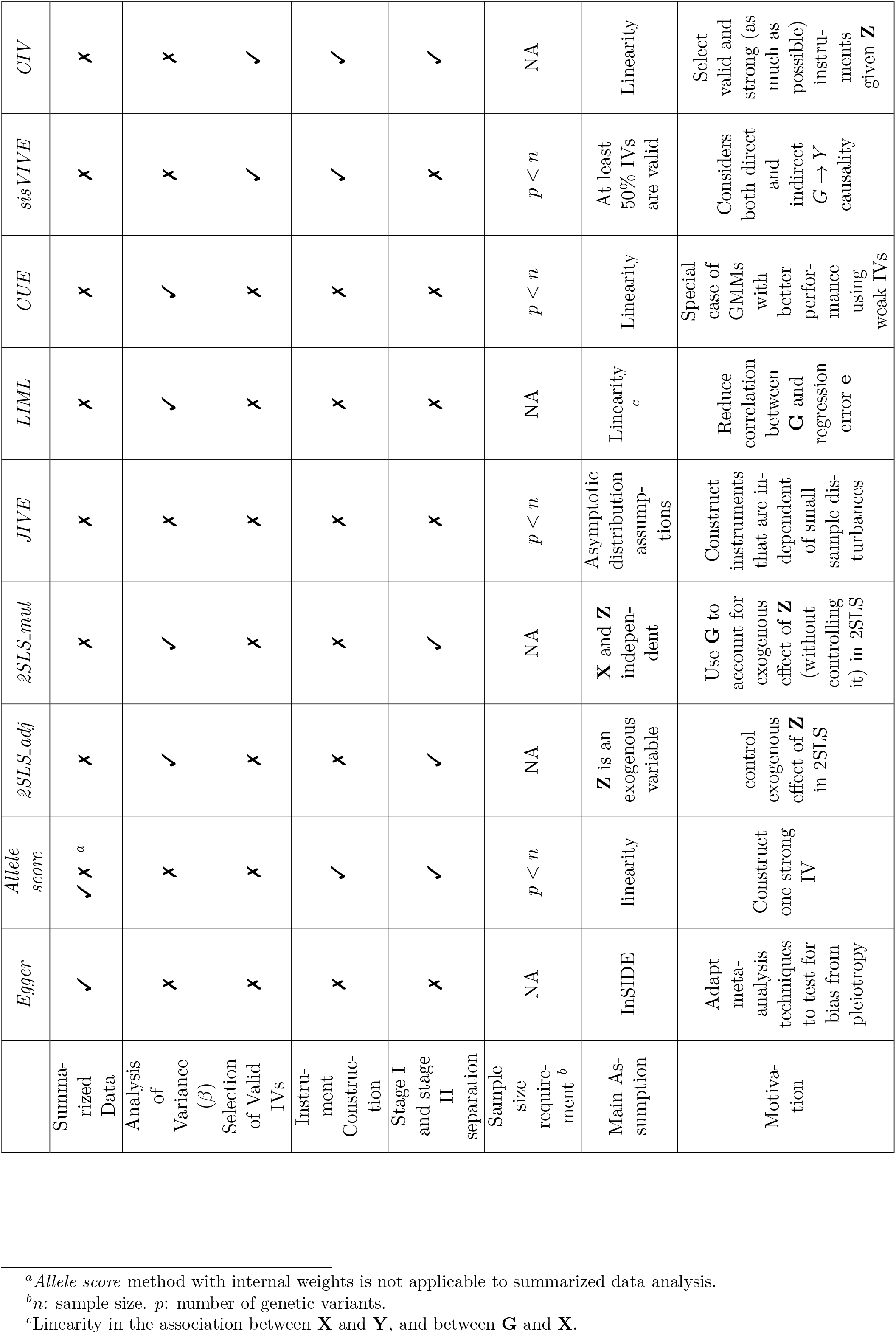
Properties of selected Mendelian rah3domization methods. Columns: Selected instrumental variable methods. Rows: Selected properties. ✓: Yes; the corresponding method has this property, or can be used in this scenario. ✗: No.

## 3 Constrained Instrumental Variable method

The Constrained Instrumental Variable (*CIV*) method proposed here is designed to maximize instrument strength yet provide robustness to pleiotropic effects, specifically for the situation where potentially pleiotropic phenotypes (**Z**) are measured and available.

### 3.1 CIV when p < n

Specifically, we are interested in finding a weight vector 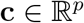 and 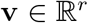, s.t.

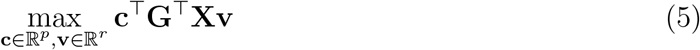

subject to conditions:

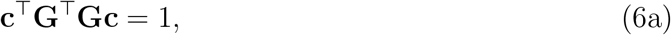

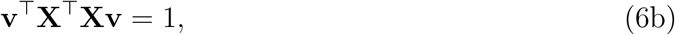

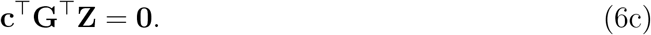

The motivation of this method is to construct a relatively strong instrumental variable **Gc** (a linear combination of variables in **G**) that is uncorrelated with **Z**. In this way the weak instrument bias and the pleiotropic effect are alleviated in Mendelian randomization with the new instrumental variable. We use Equation (5) to obtain the maximized canonical correlation between **Gc** and **X**. In addition we use Equation (**6c**) to force the new instrumental variable **Gc** to be orthogonal to all possible pleiotropic phenotypes in **Z**. This maximization problem is well-defined when *p* ≥ *k* (see Appendix A).

The strength of the *CIV* instruments can be measured with an F-statistic or with a concentration parameter (Stock et al., 2012). The former can be calculated as the F-statistic from a linear regression model against the null that the excluded instruments are irrelevant in the first-stage regression **X** ~ **G**. The latter measures the overall association between **X** and **G** without considering the number of instruments used. As a rule of thumb, a single instrumental variable with F-statistics < 10 is usually considered weak instrument. *CIV* is designed to retain instrument strength, however, it may not always yield the strongest global F-statistic since the linear constraint (6c) may force our method to exclude some genotypes, that are associated with both **X** and **Z**, from the construction of CIV.

In the context of one-sample analysis, the solution, c, is used to construct a new instrumental variable **G**\* = **Gc**. The new *CIV* is used directly to infer the causal effect of **X** on **Y** from (**G**\*, **X**, **Y**) using estimation methods for linear structural equation modeling methods such as 2SLS. Alternatively *CIV* can be embedded inside a bootstrap to find a bias-corrected *CIV* weight. We refer to the latter as “CIV_boot”.

In the context of model assessment, or when we have two different sets of participants available, we may want to split data into two datasets. Specifically, *CIV* is trained on the first (training) set, and the solution **c** is then applied to the second dataset to construct new instrumental variable **Gc** to infer causal effect. Some other methods, such as *Allele score* method can also be implemented in this way since both of the methods rely on the first stage pathway (**X** ~ **G**) to construct instruments, and this process is separated from second stage regression (**Y** ~ **X**).

One limitation of the *CIV* method lies in the fact that a solution **c** only exists when *p* < *n*. In fact, when *p* < *n*, there is an unique solution (see Appendix A). This weighted score can then be used directly to estimate the causal effect of **X** on **Y**. However, a solution only exists when *p* < *n* since **G**^⊤^**G** is not invertible when *p* > *n*. In addition, we may want to select a subset of valid instruments from among all SNPs (the columns of **G**) of interest. Therefore, we propose an extension of the previous solution that addresses these two concerns, through imposing a penalty on the problem (5).

### 3.2 Smoothed CIV

Different choices of penalty functions on the problem (5) lead to different solutions. However, popular LASSO and *L*_2_ penalties would not result in a sparse solution here under any level of regularization because of the linear constraint (6c). An explanation can be understood by examining Figure 4. The LASSO and *L*_2_ contours will touch the linear constraints (straight line in the figure) at two non-sparse solutions of c.

**Figure 4:**
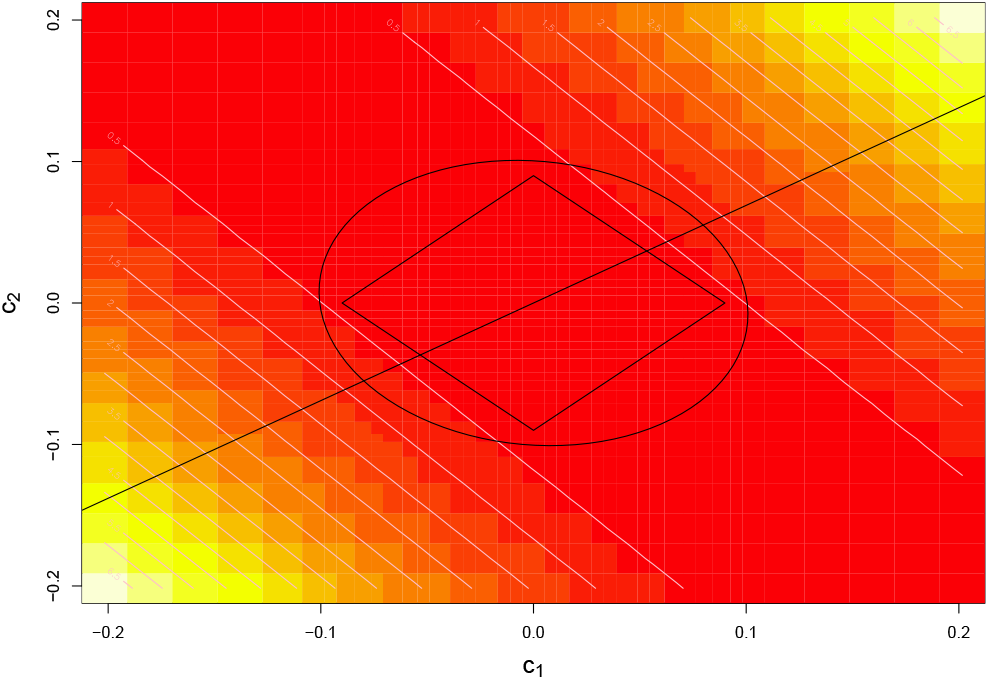
Graph demonstrating the maximization problem with LASSO penalty and *L*_2_ penalty. Rectangle: LASSO penalty contour with the same level of penalization. Circle: *L*_2_ penalty contour with the same level of penalization. Straight line represents the CIV solution space required, and it does not intersect with a sparse solution. Pixels with color from yellow to red: coordinates of **c** = (*c*_1_, *c*_2_) with absolution correlation values from high levels to low levels.

Instead we consider an *L*_0_ penalty and maximize the function

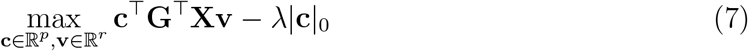

subject to conditions:

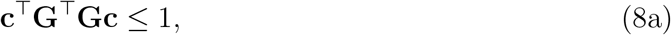

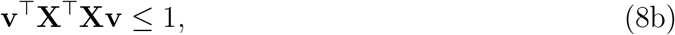

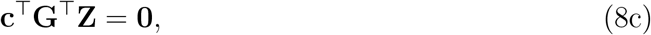

where |**c**|_0_ is the *L*_0_ norm of **c** and λ is a regularization parameter.

This problem is equivalent to maximizing a convex function over a convex set. However, it is computationally impractical to exhaustively enumerate all possible sets of |**c**|_0_; this problem with *L*_0_ norm has been proven to be NP-hard (Natarajan, 1995). Therefore, we propose instead to consider smoothed *L*_0_ penalties: 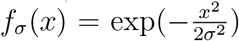. In the limit when 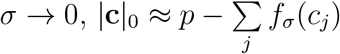, thereby the problem (7) can be approximated by:

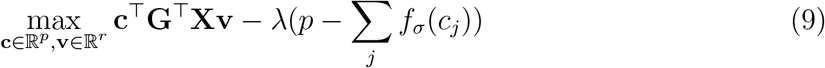

subject to conditions (8a), (8b) and (8c). The approach is then to solve problem (9) for a decreasing sequence of *σ* (→ 0) and a given value λ, while resulting in at least approximately sparse solutions. However, there are no theoretical guarantees for the uniqueness of such numerical solutions, and often there are multiple solutions.

In order to implement this smoothed L0 algorithm and obtain a single solution **c** for a given value of λ, we proceed as follows:

1. Initialization: For a given value of λ, start from an initial guess 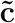 and initial *L*_0_ penalty 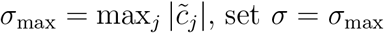.
2. While *σ* > *σ*_min_ = 0.01 we do

i. Calculate the gradient of function (7) 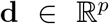, where 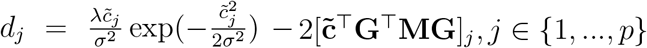 and 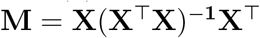.
ii. Set 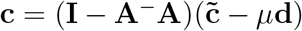 where **A**^−^ is a general inverse of **A** = **Z**^⊤^**G** and *μ* is a step-size parameter in gradient descent algorithm.
iii. Set 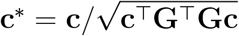 as the updated solution.
iv. Repeat (i) (ii) and (iii) (maximum *T* times) until it converges, i.e. 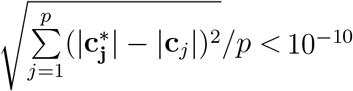.
3. Update *σ* with *σ* = 0.5 *σ*_prev_, where *σ*_prev_ is the previous value of *σ* used in step 2. If *σ* > *σ*_min_ repeat all items in step 2. If not, stop the algorithm and record the last iteration of **c** as the final solution.

In summary, the maximization problem of Equation (9) is solved by repeatedly taking gradient descent steps (i), and then projecting the possible solution back into constrained set ((ii),(iii)). Note that the step (ii) restricts the solution to be on the constrained set (8c) and step (iii) restricts to the boundary of the constrained set (8a). Note that the unconstrained gradient descent step followed by projection to the feasible set is equivalent to a direct gradient descent step on the feasible set (Cui et al., 2010). The parameters for step-size (*μ*) and number of iterations (*T*) should be carefully chosen to achieve balance between computation cost and precision. That is, the states discovered by this algorithm may not achieve the maximized value of Equation (7) even with a large number of iterations, if we use a step size that is too large. The decreasing list of values for *σ* is chosen to ensure that the approximation accuracy will gradually increase.

The tuning parameter λ affects the prediction performance of our CIV method, as for any penalization method. Higher values of λ lead to stronger penalization on the *L*_0_ norm of **c** and an approximately sparser solution for c. This means that more valid instruments will be considered as invalid and omitted from construction of the CIV weighted scores. Smaller values of λ correspond to less regularization and thus lead to less sparse solutions. The ideal value for λ will depend on the actual proportion of invalid instruments among all instruments.

We therefore implement a K-fold cross-validation technique to find an optimal value for λ that minimizes the projected prediction error 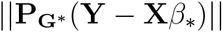, where 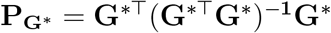. We choose to use the projected prediction error as the tuning measurement in order to make **G**\* = **Gc** as “valid” as possible rather than as “informative” as possible. In the ideal case where **G**\* is a valid instrument whose causal impact on **Y** only goes through **X**, then the regression residual **Y** − **X***β*^*^ should be orthogonal to any vectors spanned by the original instruments in **G**. In other words, only valid instruments **G** will yield 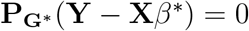. In general, we are more interested in the validity of the prediction model rather than the most informative solution of *β*; the latter may lead to over-estimation of the causal effect of interest.

There may be multiple local solutions of **c** to the smoothed problem Equation (9), since this a non-convex optimization problem. A careful reader may recognize that it implies maximization of a convex function over a convex set, yet overall this is not a convex problem! As a result, a local maximum solution of **c** may not be the global maximum solution, and numerical optimization techniques may get trapped into a local minimum. Therefore, we start from multiple (e.g. 100) initial points randomly sampled from a multivariate normal distribution *N*(0, *I_p_*), and let the smoothed *L*_0_ algorithm converges to a set of solutions, possibly arriving at multiple local modes of c. After examining correlations between all pairs of solutions (c^(1)^, c^(2)^), highly correlated solutions (corr ≥ 0.9) are removed. The remaining solutions are combined into a matrix, c*, of row dimension *p*. Finally, we construct new instruments **G**\* = **Gc**\* and refer to this approach as *CIV_smooth*.

### 3.3 Causal Effect Estimation

The causal effect estimate of exposure on response *β* is now obtained with valid instruments **G**\*. Several alternative methods can be used to infer causal effects from **G**\*, **X**, **Y**. For simplicity, we choose the two stage least square (2SLS) estimator as the default causal effect estimator. Remember that the 2SLS estimator is defined as 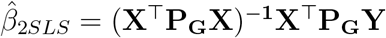; this is equivalent to implementing the least square regression twice, in the first stage **X** ~ **G**\* and the second stage **Y** ~ **X**\*. The asymptotic variance of the 2SLS estimator *β* can be estimated with:

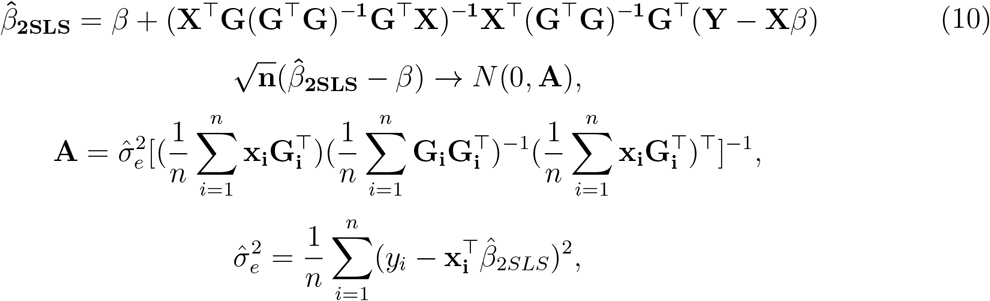

It is worth noting that **Y** − **X***β* is the regression error term which is assumed with homoscedastic variance. However, such asymptotic estimation of variance is not available for the CIV methods, since the new instruments **G**\* are dependent on all observations of **X** and **Z**, and hence 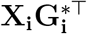 and 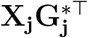 are not independent. Therefore, the assumption of weak law of large numbers is violated and puts the convergence of 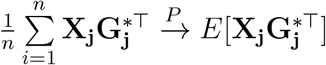 into jeopardy.

Bootstrapping can be used to estimate the sample variance of 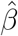 and obtain confidence intervals. The resulting empirical confidence intervals should be interpreted with caution and compared with other methods (such as the *Allele score* method) cautiously, especially when weak instruments are present (Moreira et al., 2009).

Both CIV and *CIV_smooth* can be applied to the case when two separate samples are available. For CIV construction, the weight c can be estimated on the first sample and then applied to the second sample for causal effect estimation. The same approach can be applied to the *CIV_smooth* algorithm. An alternative way to apply *CIV_smooth* to the two-sample MR analysis, similar to *sisVIVE*, is to conduct valid instrument selection using the first sample with *CIV_smooth*, and then to conduct causal effect estimation with that information on a second sample. All these three methods are included in our two-sample analysis below.

## 4 Simulation

### 4.1 Simulation Design

Simulations have been conducted under a variety of different kinds of violations of the MR assumptions in order to compare the performance of our CIV methods with other popular methods. In all simulations, we assume that there is a fairly large set of potential genetic instruments G, and that individual level data are available for the phenotype of interest **X**, potential pleiotropic phenotype **Z**, and the response **Y**.

Table 2 provides the broad goals behind four series of simulations designed to address different types of violations of the key assumptions. All simulations assume the presence of pleiotropy, and hence assumption (3) of the MR assumptions is always violated. In Series **I**, we generate strong but pleiotropic instruments **G** for phenotype of interest **X**; this violates the assumption (A2). In Series II, we investigate the association between **X** and **Z**, by varying the direction and strength of the simulated causal relationships between these two sets of variables. In Series III, we simulate a set of weak, pleiotropic instruments, where the associations between **G** and **X** are not strong; this violates assumption (A2) and jeopardizes (A1) of the three MR assumptions. Finally, in Series IV, we examine performance when the selection of important SNPs is desirable, such as what one might expect if many variants of interest are simultaneously considered within an MR analysis. Parameter settings in common across scenarios are given in Table 2. It is worth noting that we do not include simulations with *p* ≥ *n*, i.e. when there are more genotypes in **G** than observations. The reason is *CIV* and *CIV_smooth* are the only methods that would work under such restrictions and we do not have competitors in this case.

**Table 2:** Some of the parameter settings used in the four series of simulations. *n*: number of individuals. *p*: number of genotypes in total. *p_z_*: number of pleiotropic genotypes with no effect on **X**. MAF: minor allele frequency of all SNPs in the simulation. Methods: the methods compared in each simulation in one-sample setups. *α_x_*: association parameter between **G** and **X** as in Figure 3. *α_z_*: association parameter between **G** and **Z** as in Figure 3. *F_x_*: the expected F-statistic values for testing the strength of instruments in **G** (for **X**) given values of *α_x_, α_z_* and other parameters. All methods used in one sample analysis: 2SLS_naive, 2SLS_adj, 2SLS-mul, JIVE, Allele Score, *Egger* Regression, *LIML, sisVIVE*_adj, *sisVIVE*_exo, *CUE, CIV, CIV_smooth* and *CIV_smooth.sel*.

| Simulation Series | n | $\eta$ | $p$ | $p_z$ | MAF | $\alpha_x$ | $\alpha_z$ | $F_x$ | Methods (One Sample) |
| --- | --- | --- | --- | --- | --- | --- | --- | --- | --- |
| I. Standard Pleiotropy | 500 | 1.0 | 9 | 2 | 0.33 | 1 | 1 | 42.86 | All methods |
| II. $\mathbf{X} \rightarrow \mathbf{Z}$<br>or $\mathbf{Z} \rightarrow \mathbf{X}$ | 500 | 1.0 | 50 | 15 | 0.33 | 0.1 ; 0.5 | 0.1 ; 0.5 | 3.6;10.04<br>3.57;9.75 | All methods |
| III. Weak Instruments | 500 | 1.0 | 9<br>25<br>100 | 2<br>5<br>22 | 0.33 | 0.2,0.5<br>0.12,0.3<br>0.06,0.15 | 0.2,0.5<br>0.13,0.32<br>0.06,0.15 | 6.64,26.42<br>3.22,8.84<br>0.8,2.26 | All methods |
| IV. Selection of Valid Instruments | 500 | 1.0 | 50 | 10 | 0.33 | 2 ; 0.1 | 0.5 | 11.13; 2.44 | *CIV Variants*<br>*sisVIVE Variants* |

Table 3 summarizes the implementation details of the specific methods used for causal inference in our simulations. Note that we have two variants of *CIV* and *CIV_smooth* methods: *CIVJ_boot* and *CIV_smooth.sel*. The former is a bootstrapped version of *CIV*; that is, a bootstrap corrected estimate of **c** is obtained and used to infer a causal effect from (**G, X, Y**). The latter is a selection method based on *CIV_smooth*: we first obtain *CIV_smooth* estimates **ĉ**. For each converged solution **ĉ**, a feature *j* is recognized as significant if coefficient |*β*j | ≥ *ψ* *max_*j*_|*c_j_*|, *j* = 1, …, *p*. This criteria *ψ* can be tuned with cross-validation to achieve optimal selection performance in application. In our simulations we choose a small value *ψ* = 0.2 for simplicity. All selected features are then recognized as valid instruments **G^*^** for MR analyses. *CIVJ_boot* method is included in all simulations to check the consistency of *CIV*, while *CIV_smooth.sel* method is only applied in simulation Series IV to test the feature selection performance of *CIV_smooth* solutions. The description of all other methods can be found in Section 2. In series I, II and III, all methods in Table 3 are implemented for one-sample analysis, and a few comparable methods are used for two-sample analysis. In series IV, only applicable feature selection methods are considered in simulation.

**Table 3:** The methods and parameter settings used in the simulations. Label: the actual label used in figures. Parameter Choice: the parameter specifications for each of the methods in the simulations. Treatment of **Z**: the way to incorporate pleiotropic phenotype **Z** in different methods.

| Method | Label/Variants | Parameter Choice | Treatment of $\mathbf{Z}$ |
| --- | --- | --- | --- |
| 2SLS | 2SLS_naive | NA | NA |
| | 2SLS_adj | NA | Adjust $\mathbf{G} \sim \mathbf{Z}$ |
| | 2SLS_mul | NA | Combine $(\mathbf{X}, \mathbf{Z})$ to be $\mathbf{X}^*$ |
| JIVE | JIVE | NA | Combine $(\mathbf{X}, \mathbf{Z})$ to be $\mathbf{X}^*$ |
| Allele | Allele | 10 fold cross-validation | Combine $(\mathbf{X}, \mathbf{Z})$ to be $\mathbf{X}^*$ |
| | Allele_sim | “precise weight” with standard deviation 0.01 (Burgess and Thompson, 2013) | Combine $(\mathbf{X}, \mathbf{Z})$ to be $\mathbf{X}^*$ |
| Egger | Egger | NA | NA |
| LIML | LIML | NA | NA |
| | LIML_exo | NA | Adjust $(\mathbf{G}, \mathbf{X}, \mathbf{Y}) \sim \mathbf{Z}$ |
| sisVIVE | sisVIVE_adj | 10 fold cross-validation;<br>A numeric vector of penalization parameter must be given for cross-validation. | Adjust $\mathbf{G} \sim \mathbf{Z}$ |
| | sisVIVE_exo | 10 fold cross-validation;<br>list of penalization parameter | Adjust $(\mathbf{G}, \mathbf{X}, \mathbf{Y}) \sim \mathbf{Z}$ |
| CUE | CUE | NA | Combine $(\mathbf{X}, \mathbf{Z})$ to be $\mathbf{X}^*$ |
| CIV | CIV | NA | Embedded |
|  | CIV_boot | 100 bootstrap samples | Embedded |
|  | CIV_smooth | 10 fold cross-validation;<br>A numeric vector of penalization parameter must be given for cross-validation;<br>100 random initial points | Embedded |
| | CIV_smooth.sel | 10 fold cross-validation;<br>A numeric vector of penalization parameter must be given for cross-validation;<br>100 random initial points;<br>select variants whose coefficients $> 0.2 * \max(\text{coefficients})$ | Embedded |

### 4.2 Simulation Series I, II and III: Implementation

#### 4.2.1 Simulation Series I: Standard Pleiotropy

In Series I, we simulated a standard pleiotropy problem with generated values of genotypes **G**, a phenotype of interest **X**, a pleiotropic phenotype **Z** and an outcome of interest **Y**. The values of parameters were carefully chosen so that all the 9 genotypes in **G** were strong instruments for **X** and 2 of them had pleiotropic effects on **Y** via **Z**. 200 replications of the simulation for both one-sample and two-sample setups were generated. The results were compared across all approaches described in the section 2. We generated Datasets as follows:

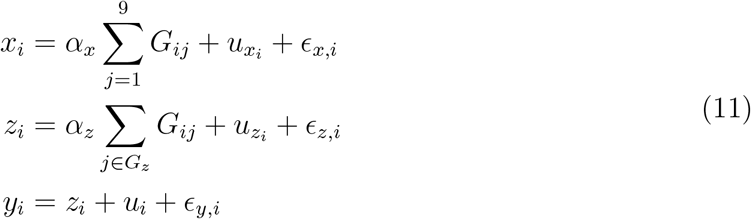

where *x_i_, z_i_, y_i_* are the ith observation of **G, Z, Y**. Let *G_ij_* denote the value of *i*th observation of *j*th genotype. *u_x_i__, u_z_i__, u_i_* represent the effect of confounding factor **U** on **X**, **Z** and **Y** respectively. *ϵ_x,i_, ϵ_z,i_, ϵ_y,i_* are the independent error of **G, Z, Y** for ith observation respectively.

The simulation of Equation (11) was based on 9 (*p* = 9) independent variants **G** simulated with a minor allele frequency 0.3 and a sample size of n = 500. The pleiotropic subset, *G_z_*, containing 2 SNPs (*p_z_* = 2) sampled without replacement, that were also associated with **Z**. Given that the SNPs were assumed to be independent with the same minor allele frequency, each SNP (coded as 0,1,2 for the number of minor alleles) had variance 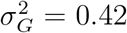. The concentration parameter, 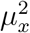, the amount of the variation in the exposure that was jointly explained by the instruments, could therefore be written as 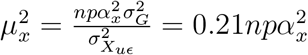, where 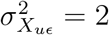 is the variance of the regression residual (*ϵ_x_* + *u_x_*) in the first stage of Equation (11). The concentration parameter divided by *p*, given the OLS estimates of *α_x_* and 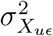, was equal to the F-statistic for testing the null hypothesis *α_x_* = 0. We set *α_x_* = *α_z_* = 1 to ensure **G** were strong instruments (F statistics > 10) for both **X** and **Z**.

We implemented simulations in both one-sample and two-sample setups for Series I, in which specific MR methods (in Table 3) were compared. The one-sample setup corresponded to a sample of n = 500 observations of **G, X, Z, Y**. In addition, we assessed performance of our *CIV* instruments and compared with appropriate comparators, including *Allele score* methods and *sisVIVE* methods in the two-sample setup. Specifically, we generated two different sets of **G, X, Z, Y**, and each set contains 500 observations. We analyzed the first dataset with the variants of *Allele score* methods, *sisVIVE* methods and *CIV* methods (see Table 3). The weights obtained from the first dataset for instrumental variable construction were then applied to the second data set. The corresponding measurements of instrument strength (F-statistic), correlations between new instruments (**G^*^**) and pleiotropic phenotype **Z**, and causal effect estimation bias from second data were recorded and compared in Figure 5. We chose the F-statistic as a measure of instrument strength instead of the concentration parameter (in one-sample analysis) because we have different numbers of instrumental variables from different methods, and only the F-statistic takes that into account.

**Figure 5:**
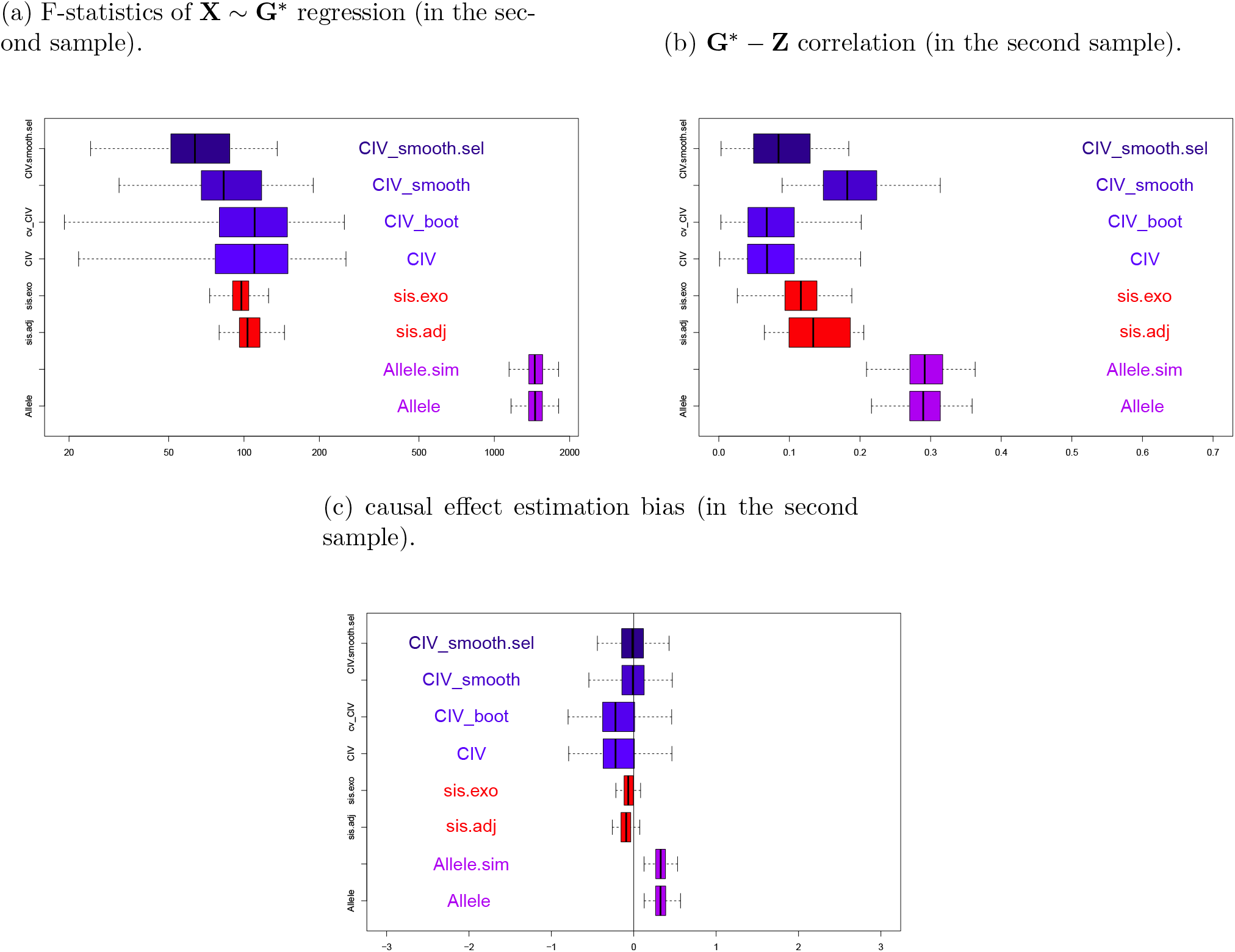
Boxplots of instrument strength measurements (a), correlation between new instruments (**G**\*) and **Z** (b), and causal effect estimation bias (c) from a two-sample set-up, with *p* = 9, *α_x_* = *α_z_* = 1 instruments across 200 simulations in series I, when true *β* = 0.

#### 4.2.2 Simulation Series II: Direct Causal Effect between X and Z

In Series II, we simulated direct causal links between **X** and **Z** to study the impact of pleiotropy on causal effect estimation of all applicable methods in both one-sample and two-sample set-ups. Specifically, **G** were generated as instruments for **X** and a proportion (*p_z_*) of them were generated as genetic causes for phenotype **Z**. Direct causal relationships between **X** and **Z**, either **X** → **Z** or **Z** → **X**, were then simulated. The outcome **Y** was depending on both of **X** and **Z**. In this way, the correlation between **X** and **Z** was induced partially through direct causal relationships and partially through overlapping genetic causes.

As far as the direct causal relations between **X** and **Z** is concerned, we find that the violation of MR assumption (A2) is inevitable when **X** → **Z**. Specifically, when there are direct causal relations **Z** → **X** and not all genetic variants in **G** are associated with Z, the situation can be considered as pleiotropy and only the genetic variants not directly related with **Z** are valid instruments. However, if the causal relation goes from **X** → Z, then all genetic variants in **G** are invalid and this should not be considered as pleiotropy. In addition, when there is a strong link between **X** and **Z**, the two sets of variables are likely to be highly correlated, and this in itself can lead to instability of the estimation of causal effect, *β*.

In Simulation Series II, datasets were generated containing direct causality between *x* and **Z** as follows:

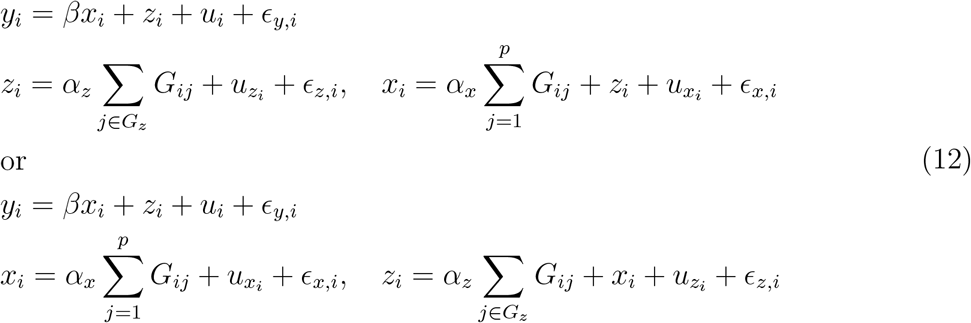

In each simulated dataset, 50 SNPs (**G** in Equation (12)) with a minor allele frequency 0.3 and *n* = 500 observations were generated. Both directions of the causality between **X** and **Z** were considered, i.e. in one case *G* → *X* → *Z* was generated, and in the other case *G* → *Z* → *X* was generated. In each case, *α_x_* ∈ (0.1, 0.5) encompassed one scenario with weak instruments and another with strong instruments. 200 datasets were generated for each scenario, and results compared the estimates and variance of the causal effect, *β*.

We ran both one-sample and two-sample simulations in Series II using the methods introduced in Table 3. The instrumental variable strength, correlation between **G^*^** and **Z**, and causal effect estimation bias in two-sample simulations of selected methods were summarized in Figure 6 and Figure 7. The performance in one-sample simulations under the causal null, when there were no true associations (*β* = 0), was reported in Figures 10 and 11 for the scenarios **X** → **Z** and **Z** → **X**.

**Figure 6:**
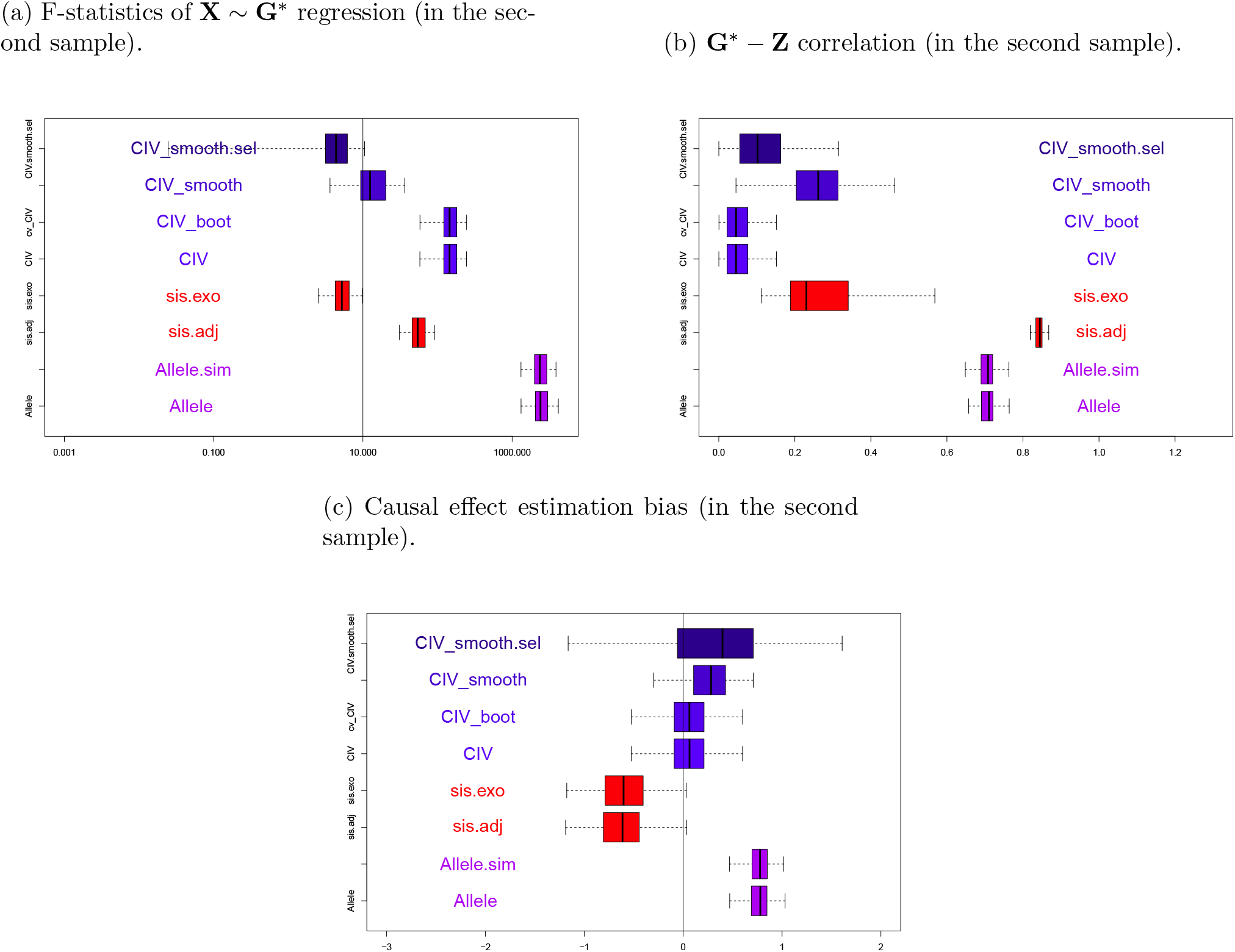
Boxplots of instrument strength measurements (a), correlation between new instruments (**G**\*) and **Z** (b), and causal effect estimation bias (c) from a two-sample set-up, with **Z** → **X** and *α_x_* = *α_z_* = 0.5 across 200 simulations in series II, when true *β* = 0. A vertical line of *F* =10 is drawn on the F-statistics plot.

#### 4.2.3 Simulation Series III: Weak Instruments

In Series III, a set of genotypes **G** (*p* = 9, 25 or 100) were generated as weak instruments for **X**. A proportion of them (2/9, 5/25 or 22/100) had pleiotropic effects on **Y** via **Z**, in which case both phenotypes (**X** and **Z**) were endogenous variables. We generated samples of (**X, Z, Y**) according to Equation 13. The choices of parameter values, as in Table 2, were based on Davies et al. (2015), and the results of all approaches (in both one-sample and two-sample setups) were displayed and discussed.

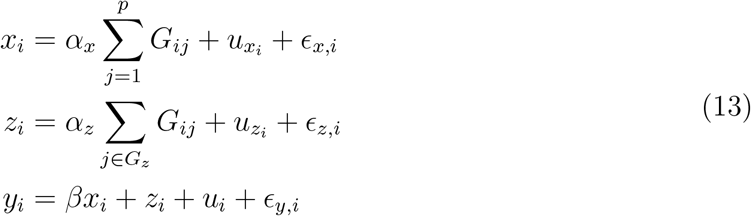

Sets of independent variants **G** in Equation (13) were generated with *p* = 9, 25, or 100, each with a minor allele frequency of 0.3 and a sample size of *n* = 500. The pleiotropic subset, *G_z_* (*p_z_* = 2, 5 or 22 respectively), containing genotypes sampled without replacement, was also associated with **Z**. The values for parameter *α_x_* were carefully chosen so that, for different numbers of SNPs *p*, all variants in **G** were weak instruments (F statistics < 10) with the same magnitude (*μ_x_*). We also chose the values for *α_z_* and *p_z_* = |*G_z_*| (number of pleiotropic genotypes) to make sure the magnitudes of the concentration parameter for the pleiotropic effect (*μ_z_*) were the same for different values of *p*.

The parameter combinations evaluated in specific simulation scenarios were recorded in Table 4, capturing simultaneously the effects of weak instruments and pleiotropic variants, with a range of different numbers of instruments (*p* = 9, 25,100). The performances of *CIV* instruments in two-sample setup were illustrated in Figure 8. Performances under the causal null, when there were no true associations (*β* = 0), in one-sample setup were summarized in Figures 12, 13 and 14, for different values of *p*. The performance of all methods under true value *β* =1 can be found in the Appendix (Figures 19, 20 and 21).

**Table 4:** Simulation scenarios for Series III simulations with weak instruments. Data were generated for each simulation scenario with two values of *β* ∈ (0,1), and replicated 200 times (i.e. 200 datasets were generated).

| | $p$ | $p_z$ | $\alpha_x$ | $\alpha_z$ | $\mu_x^2$ | $\mu_z^2$ |
| --- | --- | --- | --- | --- | --- | --- |
| Scenario1 | 9 | 2 | 0.2 | 0.2 | 37.8 | 8.4 |
| Scenario2 | 9 | 2 | 0.2 | 0.5 | 37.8 | 52.5 |
| Scenario3 | 9 | 2 | 0.5 | 0.2 | 236.25 | 8.4 |
| Scenario4 | 9 | 2 | 0.5 | 0.5 | 236.25 | 52.5 |
| Scenario5 | 25 | 5 | 0.12 | 0.13 | 37.8 | 8.4 |
| Scenario6 | 25 | 5 | 0.12 | 0.32 | 37.8 | 52.5 |
| Scenario7 | 25 | 5 | 0.3 | 0.13 | 236.25 | 8.4 |
| Scenario8 | 25 | 5 | 0.3 | 0.32 | 236.25 | 52.5 |
| Scenario9 | 100 | 22 | 0.06 | 0.06 | 37.8 | 8.4 |
| Scenario10 | 100 | 22 | 0.06 | 0.15 | 37.8 | 52.5 |
| Scenario11 | 100 | 22 | 0.15 | 0.06 | 236.25 | 8.4 |
| Scenario12 | 100 | 22 | 0.15 | 0.15 | 236.25 | 52.5 |

### 4.3 Simulation Series I, II and III: Results

#### 4.3.1 Instrument Strength

Figure 5 shows one example of the instrument strength results from the two-sample simulations in Series I. It can be seen from panel (a) in Figure 5 that all methods (Allele score methods, sisVIVE methods and *CIV* methods) form strong instrumental variables (F-statistics > 10), and *CIV* methods create instrumental variables that have weak correlation with **Z** (panel (b)). Allele score methods form the strongest instrumental variables among all competitors; however, their causal effect estimations are biased (panel (c)) because of the relatively large correlations of allele scores and **Z** (panel (b)). sisVIVE methods also lead to strong instruments (panel (a)) with unbiased causal effect estimation (panel (c)), however, the correlation with **Z** is a bit larger than *CIV* (panel (b)). In summary, *CIV* methods (including *CIV, CIV_boot* and *CIV_smooth.sel*) form instrumental variables that have reduced correlation with pleiotropic phenotype **Z**, and lead to unbiased causal effect estimates in simulation Series I.

In Figure 6 and Figure 7 where results are shown from Series II with *Z* → *X* and *X* → *Z* respectively, all the variants of *Allele* methods and *sisVIVE* methods do not perform well. There are pleiotropic correlations of allele scores with **Z** and biased causal effect estimates from *Allele* methods. The *sisVIVE.exo* method selects instrumental variables that are relatively weak, and *sisVIVE.adj* method select pleiotropic instruments. BothsisVIVE methods give slightly biased causal effect estimates. In contrast, all *CIV* instruments show lower pleiotropic correlations with **Z** than *sisVIVE* or *Allele* methods, and the resulting causal effect estimates are less biased than the *Allele* and *sisVIVE* methods.

**Figure 7:**
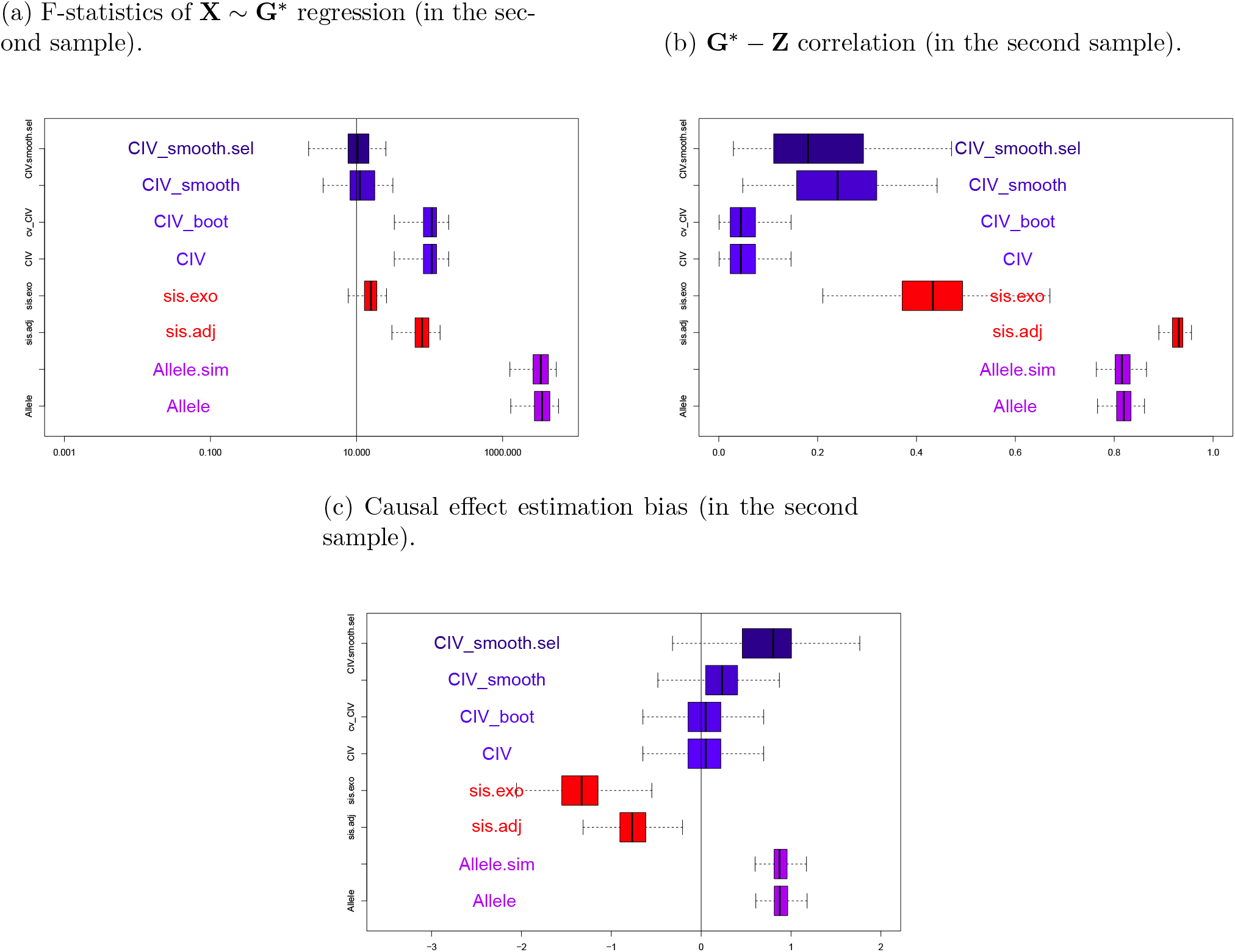
Boxplots of instrument strength measurements (a), correlation between new instruments (**G**\*) and **Z** (b), and causal effect estimation bias (c) from a two-sample set-up, with **X** → **Z** and *α_x_* = *α_z_* = 0.5 across 200 simulations in series II, when true *β* = 0. A vertical line of *F* =10 is drawn on the F-statistics plot.

However, in Figure 8 where results are shown for Scenario 9 in Series III where *p* = 100, i.e. many weak instruments, the *CIV* methods do not perform well compared to Allele score methods. It can be seen from panel (a) in Figure 8 that all *CIV* methods form relatively weak instrumental variables compared to *Allele* score methods. *CIV* and *CIV_boot* result in instrumental variables with smaller correlations with Z. Of the *CIV* methods, *CIV_smooth*, with or without selection, results in a stronger instrument with comparable strength to *sisVIVE*, and with the advantage of lower correlations with **Z** than sisVIVE, although the pleiotropic correlations with **Z** are no longer zero after smoothing of *CIV*. Here, the *Allele* methods perform extremely well, forming strong instruments that are uncorrelated with Z. It is worth noting that any weighted score based on a non-sparse weight will have a weak correlation with Z, since there are many valid but weak instruments (75/100) and the variance of **Z** is relative large in this case. Similar results were obtained for other scenarios in Series III simulations. The weak instruments from *CIV_boot* and *CIV* then result in large variability in the causal effect estimates (panel (c)). In conclusion, one limitation of *CIV* methods lies in the fact that the strength of *CIV* instruments may be excessively reduced when there are many weak instruments present in **G** (*F* < 10).

**Figure 8:**
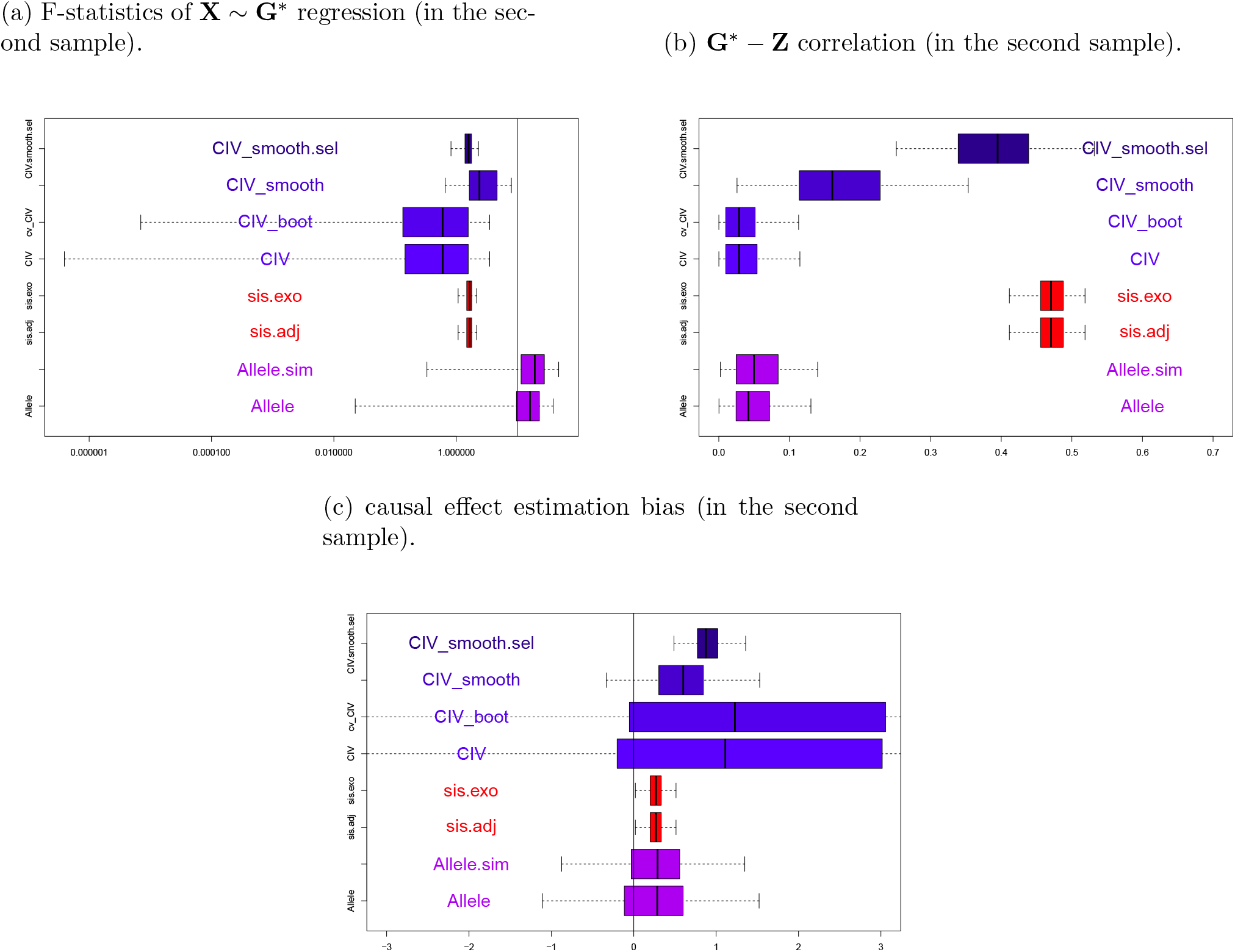
Boxplots of instrument strength measurements (a), correlation between new instruments (**G**\*) and **Z** (b), and causal effect estimation bias (c) from a two-sample set-up, with *p* = 100, *α_x_* = 0.06, *α_z_* = 0.06 instruments across 200 simulations in series III, when true *β* = 0. A vertical line of *F* = 10 is drawn on the F-statistics plot.

#### 4.3.2 Causal Effect Estimation

Figure 9 shows causal effect estimation results from Series I in a one sample set-up. It can be seen that the estimates from *2SLS-_adj* have consistent negative bias and *2SLS-naive* have positive bias, while *LIML* methods seem to show either a positive or negative bias, depending on whether or not adjustments for **Z** are included. The causal effect estimates from *sisVIVE* and *Egger* methods are slightly biased, while *JIVE* and *Egger* methods have large variability. *CUE, Allele* and *2SLS_mul* have consistently unbiased causal effect estimates and little variability across replications. All 3 flavors of *CIV*, especially *CIV_smooth*, also perform well with little bias and variability that is comparable to that of *2SLS_mul, Allele* or *CUE*. In summary, the causal effect estimation bias is small and comparable for *2SLS_mul, Allele, sisVIVE, CUE* and all three *CIV* variants.

**Figure 9:**
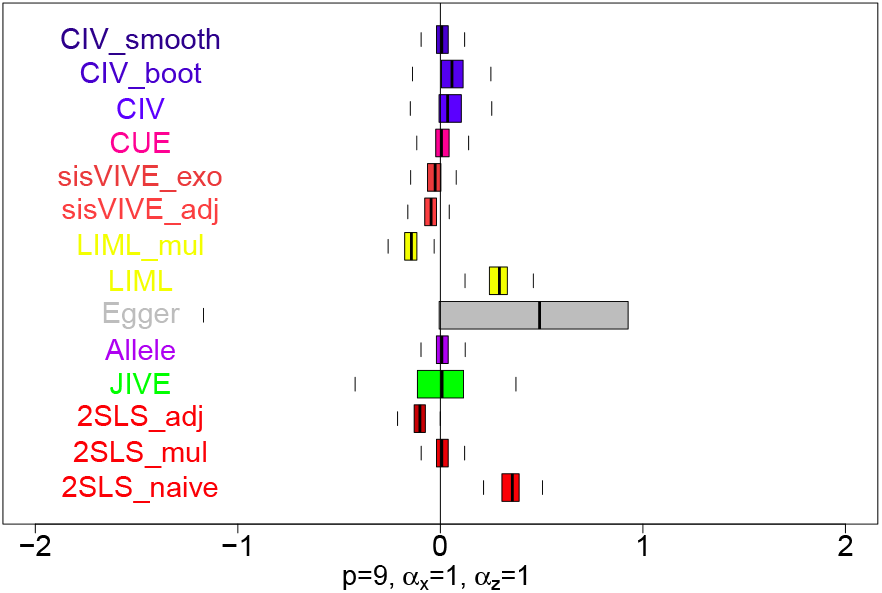
Boxplots of estimates of the causal effect estimates, *β*, from a one-sample set-up in simulation series I, with true causal effect *β* = 0 across 200 simulations.

In fact, *CIV_smooth* seems to perform the best among three *CIV* variants, with less bias and variability than the others. *CIV_smooth* method also has slightly less bias than *sisVIVE*, the reason is that *sisVIVE* does not consistently select valid instruments across replications according to our observations. Here, the regression adjustments for **Z** used by *2SLS* is biased since the adjusted instrumental variables (regression residuals) violate the assumption (A1). Note that *Allele* score method performs well here in one-sample analysis in contrast to its biased result in two-sample analysis (8). In conclusion, using the smoothed solution of *CIV*, the impact of pleiotropic genotypes can be reduced and the causal effect estimates is consistently less biased than its closest competitors *(Allele* score and *sisVIVE* methods).

The results of one-sample simulations in Series II, when there are causal relations between **Z** and **X**, are found in Figures 10 and Figure 11. In Figure 10, we have a causal pathway from **X** to **Z**. Conditioning on **Z** directly could lead to collider bias, but since some SNPs are not directly associated with **Z**, *CIV* methods may be able to alliviate the bias by removing them. In Figure 11, the causal effect estimation results of all methods when there is a causal pathway from **Z** to **X** are shown. In this case, *CIV* methods are expected to eliminate the pleiotropic bias since they are designed to remove the pleiotropic genotypes and restore the assumption (A2). In both these figures, particularly when the instruments are strong, there are large biases associated with *sisVIVE, LIML, 2SLS-uaive* and *2SLS-adj* methods. *sisVIVE* method has a particularly large bias in Figure 10. *CUE* continues to show good performance, as does *2SLSmul* method. There is little bias for *Allele, Egger*, or *JIVE* but with weak instruments, these methods are quite variable.

**Figure 10:**
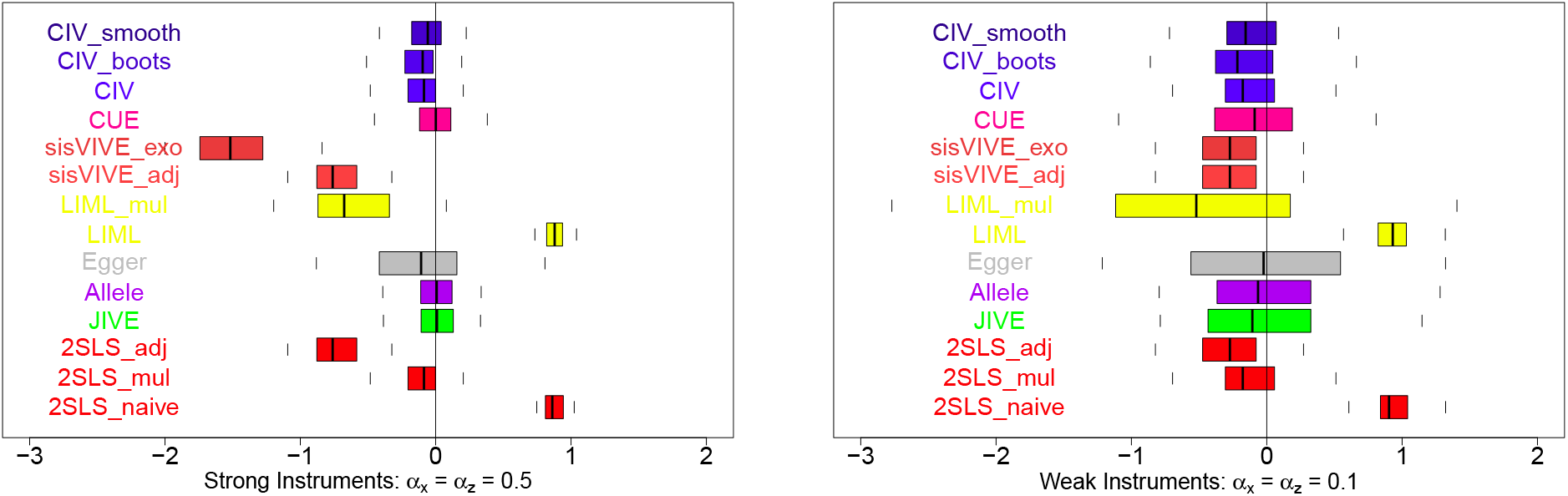
Boxplots of estimates of the causal effect estimates, *β*, from a one-sample set-up in simulation series II, when there are causal effects **X** → **Z** and *β* = 0 across 200 simulations.

**Figure 11:**
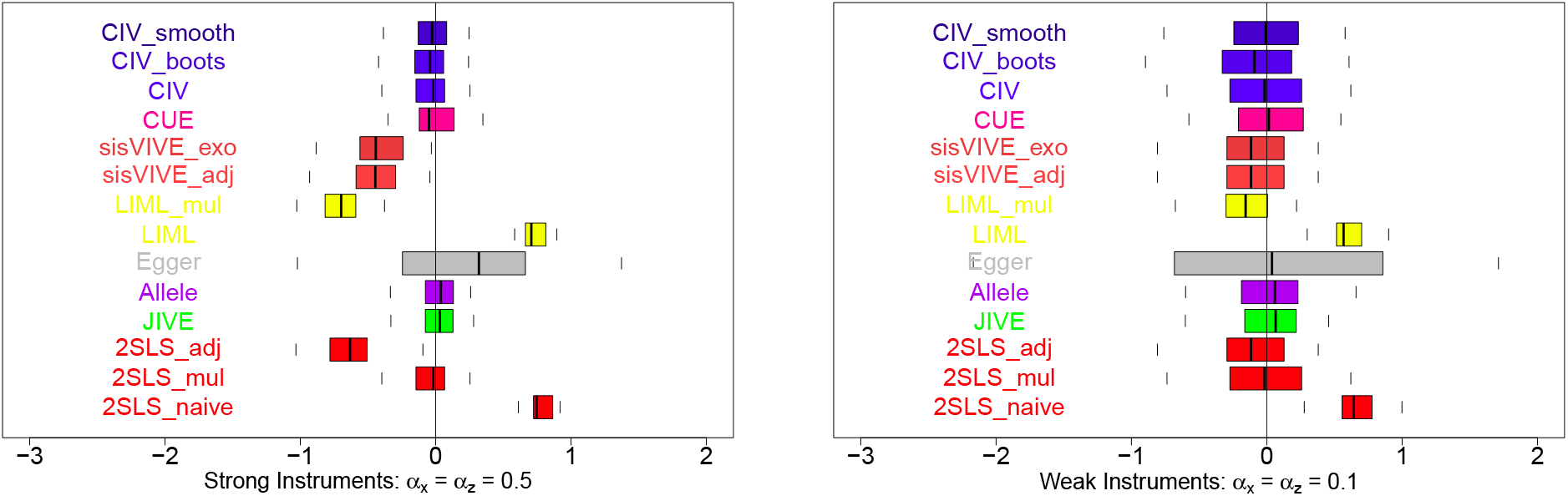
Boxplots of estimates of the causal effect estimates, *β*, from a one-sample set-up in simulation series II, when there are causal effects **Z** → **X** and *β* = 0 across 200 simulations.

All 3 flavours of *CIV* perform well here in Figures 10 and Figure 11, with little bias and variability that is comparable to that of *2SLS* or *CUE. CIV_smooth* continue to perform the best of the three *CIV* variants, with slightly less bias than the other versions.

Figures 12, 13 and 14 show causal effect estimation results from Series III, for different numbers of SNPs, *p*, in one sample set-ups. In all 3 Figures, it can be seen that the *JIVE* estimates have huge variability across simulations. *2SLS_naive* and *Egger* methods show consistent and large positive bias, and the *LIML* methods seem to show either a positive or negative bias, depending on whether or not adjustments for **Z** are included. For *CUE* and *Allele score* methods, the variability across simulations tends to increase with p. However, the variability of *CIV, sisVIVE* and *2SLS* appears to decrease as p increases.

**Figure 12:**
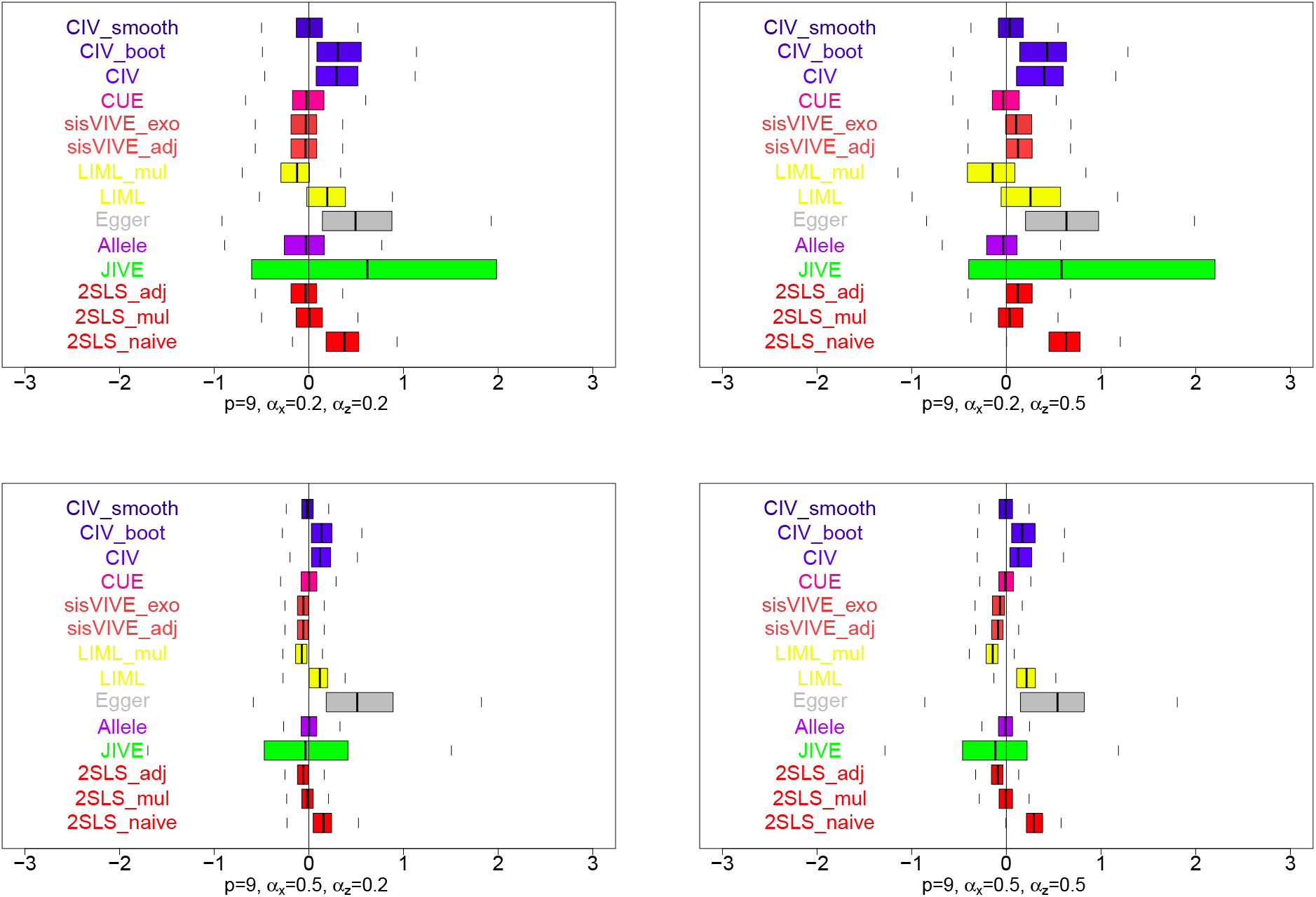
Boxplots of estimates of the causal effect estimates, *β*, from a one-sample set-up in simulation series III, with *p* = 9 instruments across 200 simulations, when true *β* = 0. The panels display results for different values of *α_x_* and *α*_z_.

**Figure 13:**
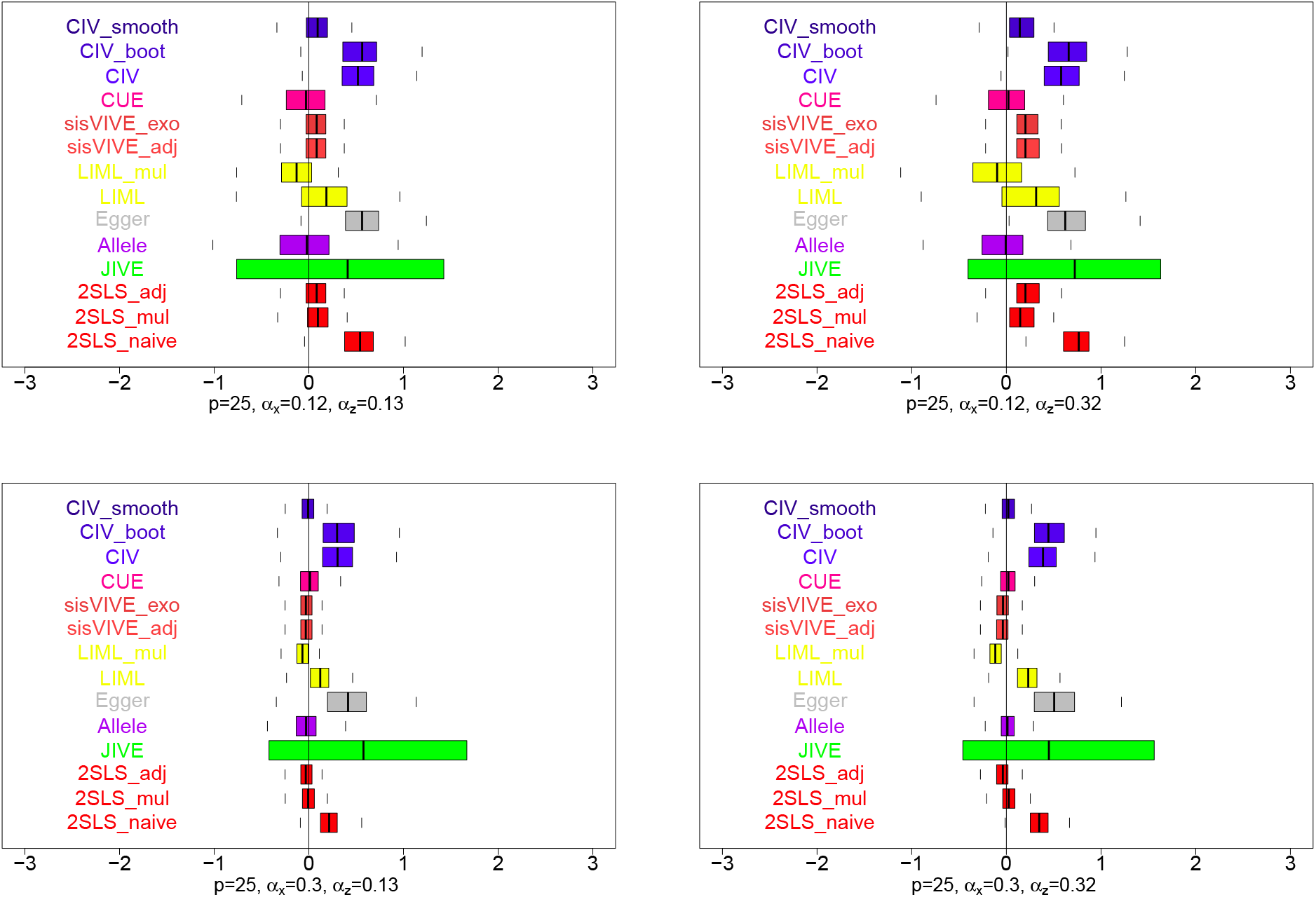
Boxplots of estimates of the causal effect estimates, *β*, from a one-sample set-up in simulation series III, with *p* = 25 instruments across 200 simulations, when true *β* = 0. The panels display results for different values of *α_x_* and *α_z_*

**Figure 14:**
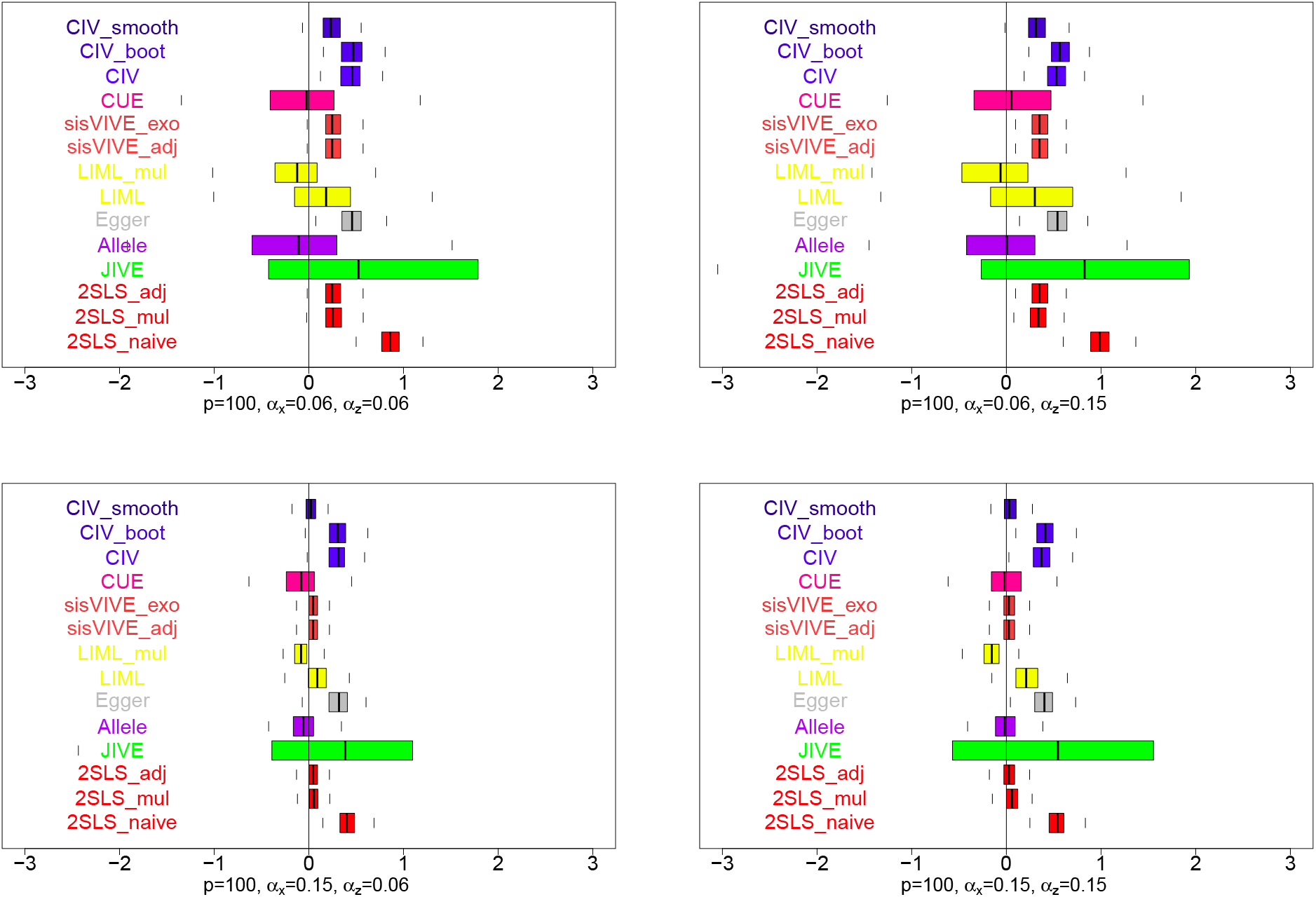
Boxplots of estimates of the causal effect estimates, *β*, from a one-sample set-up in simulation series III, with *p* = 100 instruments across 200 simulations, when true *β* = 0. The panels display results for different values of *α_x_* and *α_z_*

The only method that gives an unbiased causal effect estimate in all of 3 Figures 12–14 is *CUE*, which was explicitly designed for situations with many weak instruments. Bias is also small for *LIML* (although estimates are variable), which agrees with the general claim that generalized methods of moments are less biased than *2SLS* when using many weak instruments (Hansen et al., 2008; Davies et al., 2015).

The causal effect estimation bias is small and comparable for *2SLS_adj, 2SLS_mul, sisVIVE* and *CIV_smooth*. The similarity between *CIV_smooth* and *sisVIVE* is expected. These two methods both perform embedded feature selection, and hence are likely to underperform when only selecting a few weak instruments from a large set of candidates. The comparable performances of *2SLS_adj* and *2SLS_mul* here are probably due to the design of the simulations for Series III; there is no causal relationship between **X** and **Z** and all genotypes are weak instruments. Hence, the adjustments for **Z** used by *2SLS* work well. The biases for *Allele* score method and *CIV* without smoothing are larger; indicating that in general summarizing many weak instruments into one instrument does not provide substantial benefit here.

### 4.4 Simulation Series IV: Selection of Valid Instruments

A fourth series of simulations was designed to explore the performance of valid genotype selection in one-sample setup. Here, **G** contained not only some strong instruments (some of which may have pleiotropic effects), but also many weak instruments. We referred to these weak instruments as “unnecessary genotypes”; perhaps they were put into the pool of instruments since they shared the same genetic pathway with a strong SNP, or their contributions were so weak that they were not adding information. Use of such SNPs as potential instruments jeopardized the validity of the whole set of instruments, **G**, and causal effect estimates may be biased with some instrumental variable methods. Of particular interest here is the comparison of performance between CIV_smooth and sisVIVE, which both do genotype selection.

Data were simulated for *n* = 500 observations containing data (**G, X, Y, Z**). For each observation, *p* = 50 independent SNPs were generated with a minor allele frequency of 0.3, divided into two subsets *G*_UN_ (30 unnecessary SNPs) and *G*_R_ = *G*/*G*_UN_ (20 relevant SNPs). We also sampled 10 SNPs (as *G_R_z__*) without replacement in *G_R_* to be pleiotropic, i.e. also associated with **Z**. For each individual *i* ∈ (1,…, *n*), the phenotypes (*X_i_* and *Z_i_*) and response (*y*_i_) were simulated as described below:

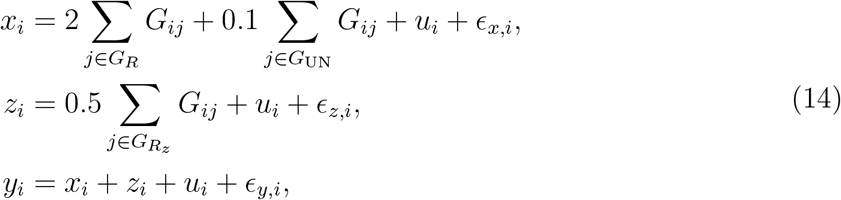

where *G_R_z__* was the subset of relevant SNPs that were associated with **Z**, and *u_i_* was a confounding factor related to **X**, **Z** and **Y**. We generated *u_i_* and *∈*. from the standard Gaussian distribution. These SNPs with *α_x_* = 0.1 were all weak instruments and not actually unrelated with *x_i_*. We used the term “unnecessary” here only because the selection of these SNPs should be largely unnecessary for the causal effect estimation purpose. We repeated the simulation 200 times and aggregated the results.

The selection of genotypes using *CIV_smooth* was conducted in two different ways. In the first approach, we selected the “top” feature, i.e. the feature corresponding to the maximum absolute magnitude of **c**, in each converged *CIV_smooth* solution. Then we combined all “top” features across different solutions and recorded the selection frequency of each feature. This method was referred as *CIV_smooth.top*. The number of features selected in this way would be limited; since many solutions only had slightly different weighting values of SNPs, but the ‘top” feature in each solution could be the same. The second selection strategy was based on *CIV_smooth.sel*. We first labeled all SNPs *j* as “relevant” in a weighting vector **c** if *C_j_* ≥ 0.2 max |c|. Then we counted the number of times each SNP j was selected, and used the averaged value across solutions as the relative selection index for all genotypes.

The results of genotype selection were shown in Table 5. In general *CIV_smooth* selected fewer of the pleiotropic genotypes in **G**, as well as fewer unnecessary genotypes. For example, *CIV_smooth* (using only the top feature) selected 3.03 valid genotypes which accounted for 52% of all selected features. This means on average 52% of the top features in all converged modes **c** corresponded to valid genotypes, while sisVIVE with either the adjusted or the exogenous **Z** methods only achieved a sensitivity below 20%. The proportion of irrelevant features and pleiotropic features were also substantially lower when using the *CIV_smooth* methods. In conclusion, the valid instrument selection performances from *CIV_smooth* were substantially better than sisVIVE methods.

**Table 5:** Simulation Series IV with unnecessary genotypes: feature selection results for CIV_smooth and sisVIVE among SNPs with different kinds of associations and across different methods. The table shows the average number (proportion) of selected features from different methods, averaging over 200 simulated data sets. *CIV_smooth.top* means only the SNP with highest |c_j_| were selected and used as instrument for MR analyses.

|  | Valid Genotypes | Pleiotropic Genotypes | Unnecessary Genotypes |
| --- | --- | --- | --- |
| *CIV_smooth.top* | 3.03 (52%) | 0.65(11%) | 2.14(36%) |
| *CIV_smooth.sel* | 7.96 (25%) | 5.47 (17%) | 18.31 (57%) |
| *sisVIVE_adj* | 10.00 (20%) | 10.00 (20%) | 30.00 (60%) |
| *sisVIVE_exo* | 6.83 (15%) | 6.25(14%) | 31.42 (70%) |

## 5 Data Analysis: Alzheimer’s Disease

Alzheimer’s disease (AD) is a chronic neurodegenerative disorder that causes a slow decline in memory and reasoning skills. It is well known that biomarkers including cerebrospinal fluid tau protein (CSF-tau) and CSF amyloid beta-protein ending at amino acid position 42 (CSF-A*β* 1-42) are reliable measures of AD progression (Iturria-Medina et al., 2016). Recently, other biomarkers such as glucose metabolism and neural functional activity have been added while exploring the mechanisms underlying late-onset Alzheimers disease (LOAD) using multi-factorial data analysis (Iturria-Medina et al., 2016). However, at this point, there is still some uncertainty whether the changes in these biomarkers “cause” AD progression or are simply associated with AD progression.

We have used instrumental variable methods on data from the Alzheimer’s Disease Neuroimaging Initiative (ADNI) (Mueller et al., 2005) to try to disentangle causal relationships for AD. ADNI began in 2004 and included 200 subjects with early AD, 400 subjects diagnosed with mild cognitive impairment (MCI), and 200 elderly control subjects. Biospecimens, including blood, urine, and cerebrospinal fluid (CSF) were collected from participants. FDG PET imaging was performed on participants within two weeks before or after the in-clinic assessments of memory composite scores. The global and regional standardized uptake value ratios (SUVR) of each subject were recorded after each scanning. Genotyping and sequencing data were also available for all subjects obtained from Illumina Human610-Quad BeadChip. Further details of of protocols and procedures of this data was available on the Image Data Archive (IDA).

The outcome (AD status) here is a binary variable, and MR methods assume linear additivity and are not designed for the situation of a binary outcome (Didelez and Sheehan, 2007). Alternatively, under more restrictive parameter assumptions (Vansteelandt et al., 2011), logistic regression and log-linear regression can be used in the second stage of MR analysis to estimate a causal risk ratio (CRR) when response is binary (Clarke and Windmeijer, 2012; Burgess et al., 2014). If only one exposure (**X**) is analyzed with one binary outcome under linear assumption in the second stage, then any bias towards the null in the causal effect estimate would be largely due to the impact of confounding factors (Palmer et al., 2008).

A very important limitation of MR analysis in ADNI data is the retrospective nature of ADNI study design. Ascertainment in ADNI was retrospective by disease status, and therefore, instruments that would be valid for a prospective study design may not remain valid after retrospective sampling in the ADNI data (Didelez and Sheehan, 2007). Specifically, the estimated first stage (**G-X**) association from case-control samples may be biased relative to the true association in a general population sample (Tapsoba et al., 2014; Tchetgen Tchetgen, 2013). If the disease being studied is rare, it is possible to use only the control samples in a first stage regression, in MR methods where two-stage methods are appropriate (Lin and Zeng, 2009). Therefore, since we realize that MR analysis of the ADNI samples is not ideal, we use this dataset simply to illustrate performance of our methods, and not to make strong causal statements. Furthermore, we only present results where the first stage associations are estimated from the controls.

### 5.1 Outcome, Exposures and Instruments

#### *Outcome* Y

A subject is either from control group or diagnosed with MCI or AD. We combine AD and MCI subjects into the group with the same response variable. We collected *n* = 491 subjects including 151 controls with outcome *Y* = 0 and 340 patients with outcome *Y* = 1.

#### *Exposures* X

We are interested in estimating the causal effect of several exposures/biomarkers including CSF amyloid beta-protein (A*β*) (*X*_1_), Phosphorylated Tau Protein (Ptau) (*X*_2_), Total Tau protein (Ttau) (*X*_3_) and FDG_SUVR (*X*_4_) on AD progression. It is well known that the isoforms of Apolipoprotein E (ApoE), a class of apolipoprotein that mediates cholesterol metabolism, have effects on both Amyloid beta aggregation and Tau protein phosphorylation, and thus there may exist pleiotropic effects in this case. Natural logarithm scales of Ttau and Ptau are used to obtain *X*_2_ and *X*_3_. All exposures (*X_k_*, k=1,…,4) (and also outcome Y) are adjusted before analysis with covariates including age, sex and education. Profiles of the subjects are summarized in Table 6.

**Table 6:** Characteristics of subjects studied in ADNI.

| | Number | Age (years)<br>(mean ± SD) | Gender(M/F) | education (years)<br>(mean ± SD) |
| --- | --- | --- | --- | --- |
| Control | 151 | 75.93 ± 5.86 | 86/65 | 16.3±2.7 |
| MCI/AD | 340 | 74.08 ± 7.63 | 212/128 | 15.89 ± 2.92 |
| MCI | 277 | 73.64 ± 7.53 | 173/104 | 16.03 ± 2.81 |
| AD | 63 | 76.03 ± 7.78 | 39/24 | 15.27 ± 3.31 |

#### *Instruments* G

For each of the exposure *X_k_*, k = 1,…, 4, the strongly associated SNPs reported by the NHGRI-EBI Catalog of published genome-wide association studies (Burdett et al., 2016) were collected from the ADNI Imputed Genotype data. The missing samples were imputed to the 1000 Genome Project utilizing the same protocol for the ROS/MAP and AddNeuroMed study. When there were very highly correlated (*ρ* ≥0.8) sets of SNPs, we kept only one representative SNP of each correlated sets. Specifically, we first built a genotype subset containing these SNPs with all pairwise correlations (*ρ* ü0.8). Then we added SNPs into the feature subset one at a time and would reject the SNP if its pairwise correlations with the selected subset were high (*ρ* ≥0.8). The final SNP set was then further reduced with univariate feature selection using significant F-statistics (*ρ* ≤ 0.05). Hence, reduced sets of SNPs containing 20 SNPs for A*β* (*X*_2_), 8 SNPs for Ptau (*X*_2_), 5 SNPs for Ttau (*X*_3_) and 25 SNPs for glucose metabolism (*X*_4_) were retained for use with Mendelian randomization methods.

### 5.2 Mendelian Randomization Analysis

The assumption (A1) of Mendelian randomization stated that the SNPs must be associated with biomarkers of interest. Strong instruments with F-statistics bigger than 10 are usually preferred in MR applications. The F-statistics for instrument strength of each biomarker here (A*β*, Ptau, Ttau, SUVR) are 11.86, 12.40, 3.89 and 5.20 respectively, indicating strong instruments only for A*β* and Ptau. We also perform Sargan test for over-identification (Baum et al., 2003) to test the MR assumption (A2) and (A3). The p-values of the Sargan test are 0.22, 0.01, 0.81 and 0.92 for X⅛, k = 1,…, 4, implying the existence of invalid instruments in **G** for Mendelian randomization for Ptau (X_2_) on AD progression (**Y**). The reason is because the selected SNPs that are strongly associated with Ptau have even stronger associations with A*β*. Hence this creates a pleiotropy problem for causal inference of A*β* and Ptau on AD progression using similar groups of genotypes. More information about the associations of instruments **G** with A*β* (X_1_) and Ptau (*β*) are shown in the figure 15.

**Figure 15:**
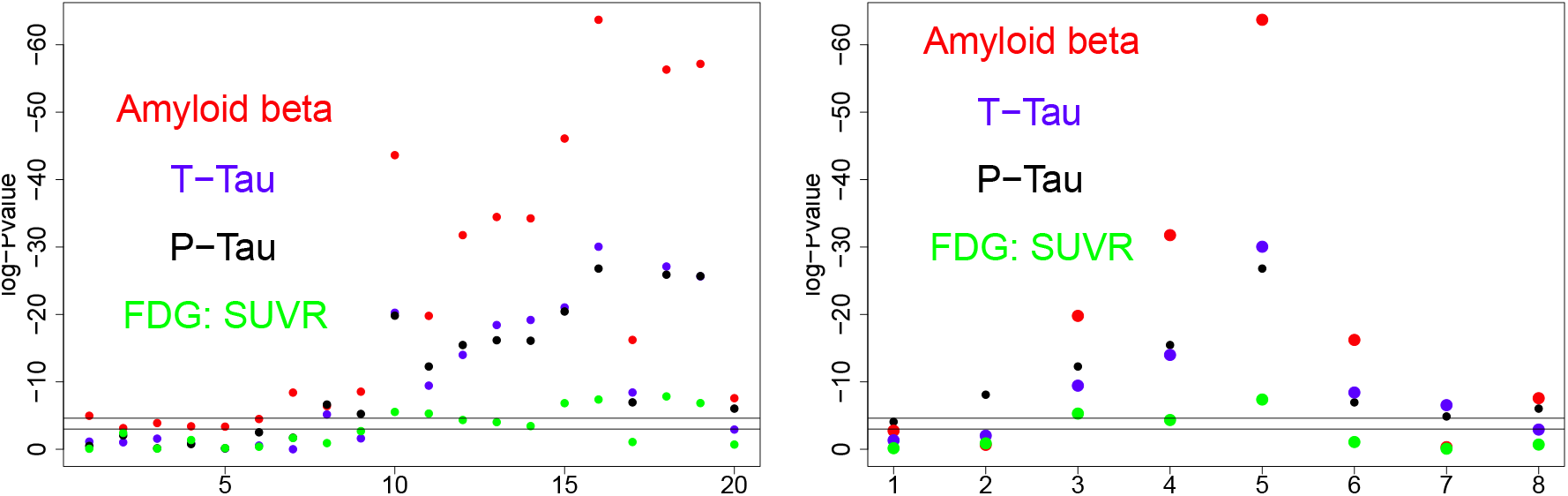
The associations between instruments selected for phenotypes and phenotypes of interest. Left: log-Pvalues for association test between the selected SNPs (for A*β*) and A*β* (*X*_1_). Right: log-Pvalues for association test between the selected SNPs (for Ptau) and Ptau (*X*_2_).

Mendelian randomization was performed to evaluate the potential causal effects of variability in all biomarkers (X) on the AD progression (Y) in a two-sample set-up. First we used only the control samples to obtain weights with the three applicable methods (*Allele, sisVIVE* and *CIV* and their variants). In the second step we constructed instrumental variables using these obtained weights with genotype information on the whole sample, and inferred causal effects of each biomarker *X_k_* on AD progression (Table 3) while treating other biomarkers as secondary phenotypes, if applicable. In this way, the retrospective nature of ADNI is respected, if we assume that the control sample is similar to the whole population from which the individuals were drawn. We can only include *CIV, Allele scores* and *sisVIVE* in the two-sample analysis because not all methods can be adapted to a two-stage approach here. Hence, the results of this MR analysis are illustrative, and require further validation, ideally on a separate prospective study.

### 5.3 Results

The 95% confidence intervals of the causal effect estimates for all four biomarkers obtained from two-sample analyses are reported in Figure 16 and Figure 17. All variants of *Allele score* methods and *CIV* methods (except *CIVJ>oot*) identified significantly negative causal effect of Amyloid beta 1-42 protein levels on AD progression. Only the *Allele score* methods and *CIV_smooth.sel* showed a significantly positive causal effect of Ptau protein levels on AD progression. Furthermore, only Allele.sim and *CIV_smooth.sel* methods identified significant causal effects of Ttau protein levels on AD progression. For glucose metabolism levels none of the instrumental variables methods showed significant causal effect estimates.

**Figure 16:**
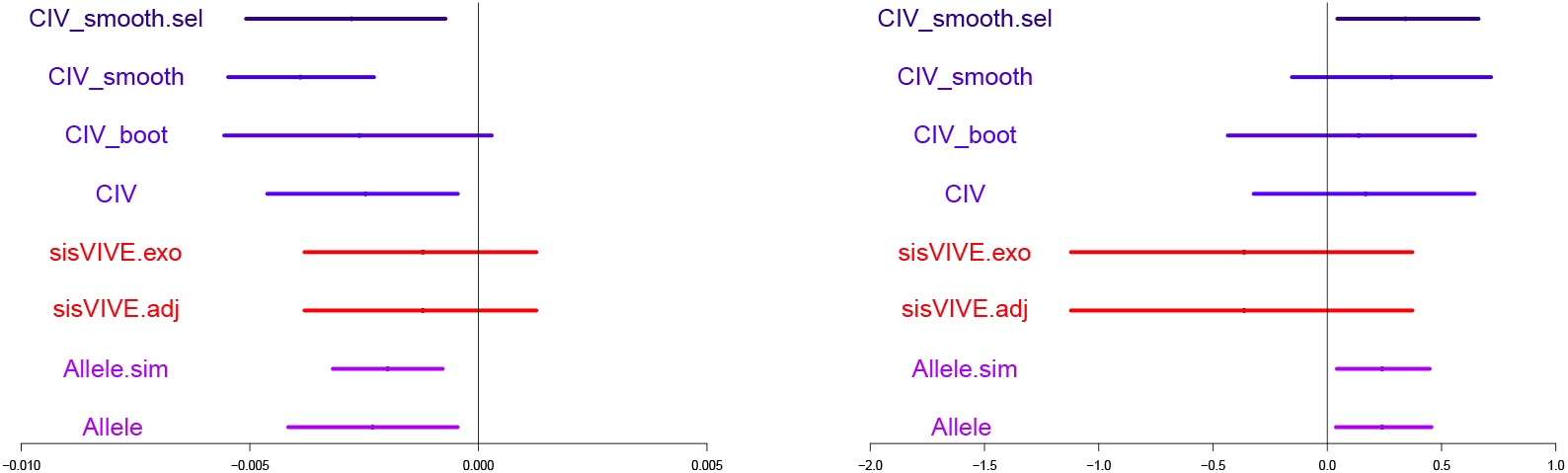
95% bootstrapped confidence interval of causal estimation of Amyloid beta 142 protein levels (supposedly negative effects) and Ptau protein levels (supposedly positive effects) on AD progression using different instrumental variable methods in a two-sample set-up. Left: Amyloid beta protein levels on AD. Right: Ptau protein levels on AD.

**Figure 17:**
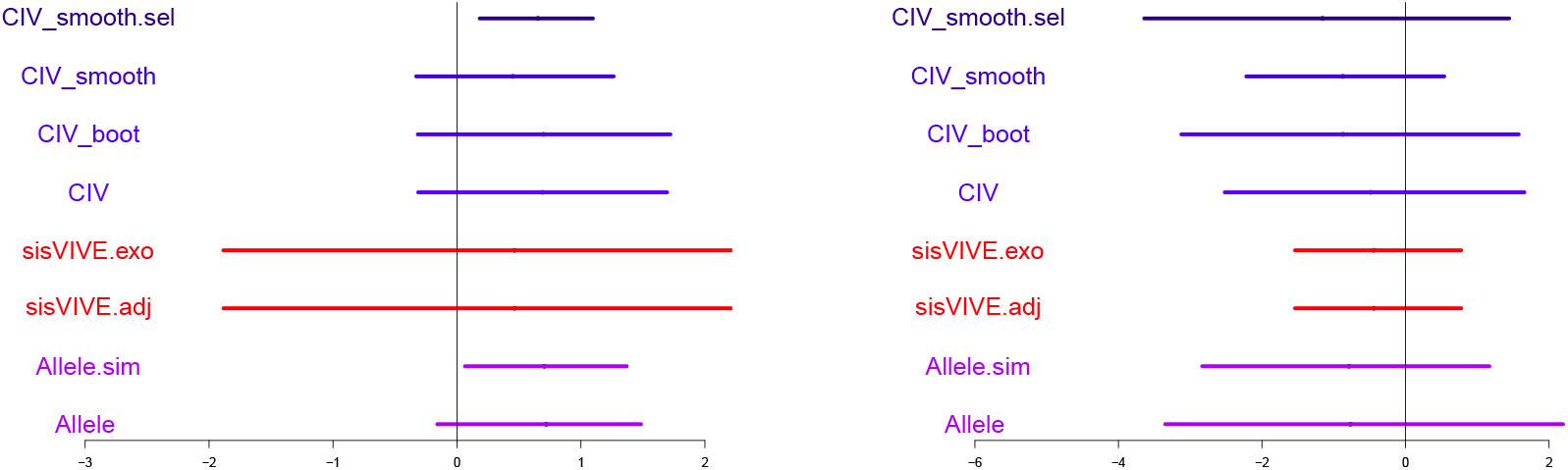
95% bootstrapped confidence interval of causal effect estimation of Ttau protein levels (supposedly positive effects) and glucose metabolism on AD progression using different instrumental variable methods in a two-sample set-up. Left: Ttau protein levels on AD. Right: glucose metabolism levels on AD.

The observation of significant causal impact for A*β* and Tau protein on Alzheimer’s disease study is consistent with some previous publications. In fact, multiple observational studies have reported decreasing A*β* and increasing Tau levels in cerebrospinal fluid of patients with Alzheimer Disease compared to normal control subjects (Sunderland et al., 2003; Maruyama et al., 2001). There is no conclusive evidence to support a significant causal relationship between glucose metabolism levels and AD progression. However, limited research has been conducted to evaluate the causal relationships of CSF A*β* proteins, Ptau, Ttau and glucose metabolism on AD using Mendelian randomization.

The range of causal conclusions from different instrumental variable methods with Amyloid beta and Ptau protein levels may be due to the existence of pleiotropy. In fact, most of the SNPs strongly associated with Ptau in this dataset were also strongly associated with Amyloid beta, and most of the associated SNPs were located in the APOE gene. As a result, some instrumental variable methods using these invalid instruments could be biased. Weighted score methods, including allele score methods and *CIV_smooth.sel*, were able to reduce bias due to pleiotropy compared to *sisVIVE*. In conclusion, when pleiotropy exists, *CIV_smooth.sel* could select valid instruments and perform adjusted causal effect estimation to account for pleiotropy.

## 6 Discussion

In this paper we proposed constrained instrumental variable methods for causal inference when pleiotropy is suspected. We have presented the performance of our methods and compared with other popular methods in different simulation scenarios (i.e. standard pleiotropy, direct causal effects between phenotypes, weak instruments, valid and invalid instruments selection). To illustrate the performance of *CIV* methods in real data, we conducted MR analysis on data from Alzheimer’s Disease Neuroimaging Initiative (ADNI) (Mueller et al., 2005) and demonstrated that the previously known associations for Amyloid beta 1-42 and Tau protein levels showed evidence for a causal relationship, even after making adjustments for other potentially-pleiotropic biomarkers.

One important feature of our *CIV* methods is that they find a balance between strength and validity in instrument construction. *CIV* solutions are designed to preserve maximum instrument strength while adjusting for the pleiotropic phenotype **Z**. In general strong instruments will provide consistent causal effect estimates, while approximately valid instruments will reduce the pleiotropy-induced bias. When the instruments are strong in a standard pleiotropy problem (Section 4.2.1), our *CIV_smooth* method yields better estimates in both one-sample and two-sample setups, and outperforms *2SLS-adj, sisVIVE* methods, *LIML* methods, *Egger* regression, and *Allele* score with less bias. In simulations with a direct causal link between **X** and **Z** (Section 4.2.2), *CIV* methods yield strong instruments while retaining less pleiotropic genotypes, compared to *sisVIVE* and Allele methods. When the instruments are weak (Section 4.2.3), our *CIV_smooth* method performs slightly worse than *CUE* and *Allele* score in terms of consistence and bias; indicating that relatively valid but weak instrumental variables may still lead to more biased results. In simulation IV (Section 4.4) where we examined valid instrument selection, we showed that *CIV_smooth* has a higher rate of valid instrument selection compared to its closest competitor, *sisVIVE*.

Another advantage of the *CIV* methods is the separation of instrument construction and causal effect estimation. In fact, the construction of *CIV* only relies on the first stage pathway **G** → **X**. Then, any estimation method for linear structural equation modeling can be applied to CIV_smooth instruments **G^*^ → X → Y** for causal inference. Due to this separation of first-stage and second-stage analysis, *CIV* (IV_smooth) and *Allele scores* can be trained and assessed on different datasets. In conclusion, *CIV* methods have substantial flexibility in terms of the model assessment and implementation of estimation methods.

Multiple solutions can be obtained with *CIV_smooth-i.e*. the solution to the problem (7) may not be unique. An example of this potential multi-modal problem is shown in Figure 18, where one simulated dataset from Series II was analyzed. The hierarchical cluster dendrogram shows the solution space for this simulation, demonstrating that there exist multiple different *CIV_smooth* solutions. However, a principal component analysis of the converged solutions, across simulations, shows that in many simulations only 1 unique solution stands out. So although multiple distinct solutions do occur, often they are extremely similar to each other. To obtain our *CIV_smooth* estimator, we attempt to sample the possible solution space by starting our converging iterations from multiple initial points, and combining all converged distinct solutions into a matrix c. This approach provides a spectrum of the possible instruments that achieve the maximized association and low correlation with **Z**.

**Figure 18:**
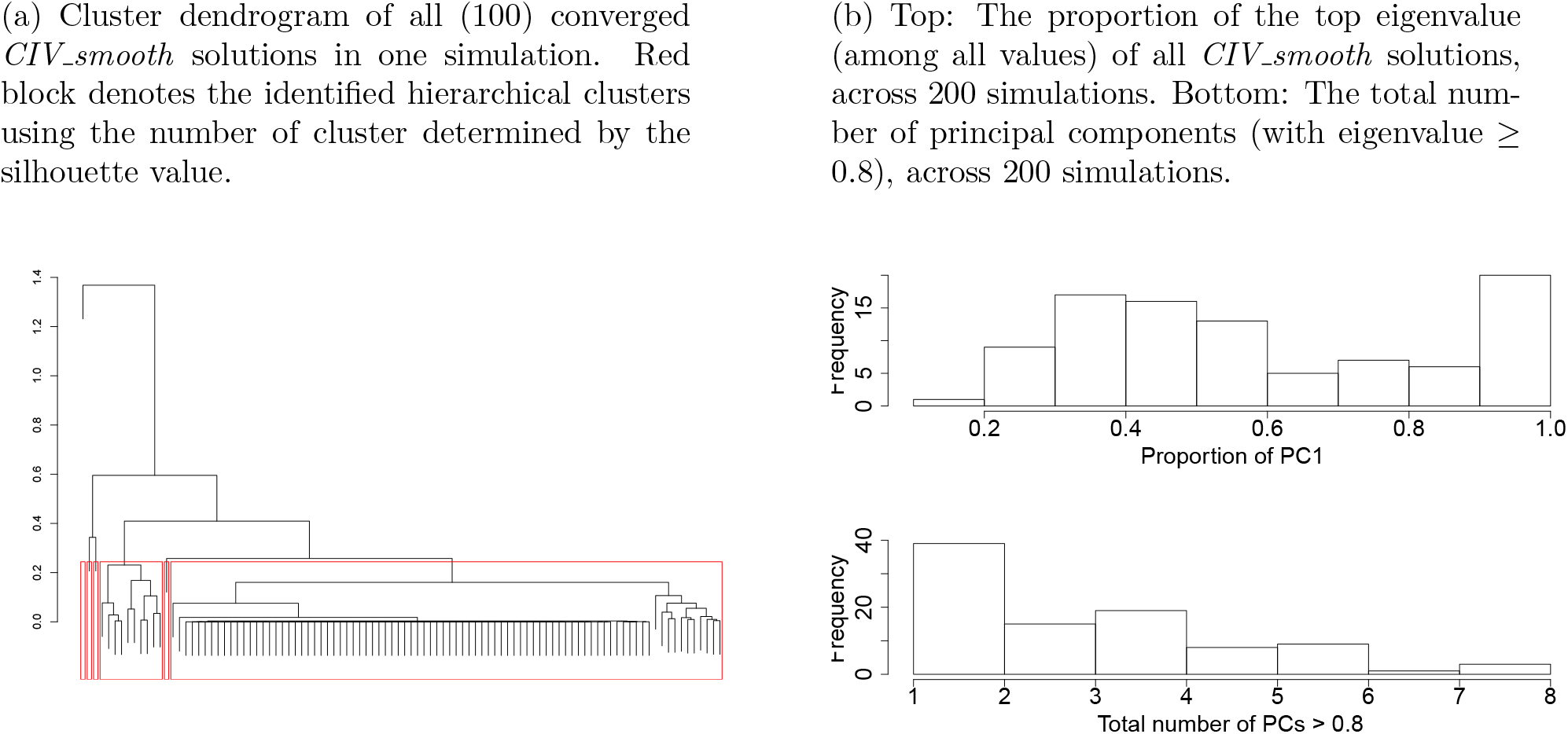
Cluster dendrogram (a) and principal component analysis (b) from a one-sample set-up, with **Z → X** and *α_x_* = *α_z_* = 0.1 across 200 simulations in Section 4.2.2.

**Figure 19:**
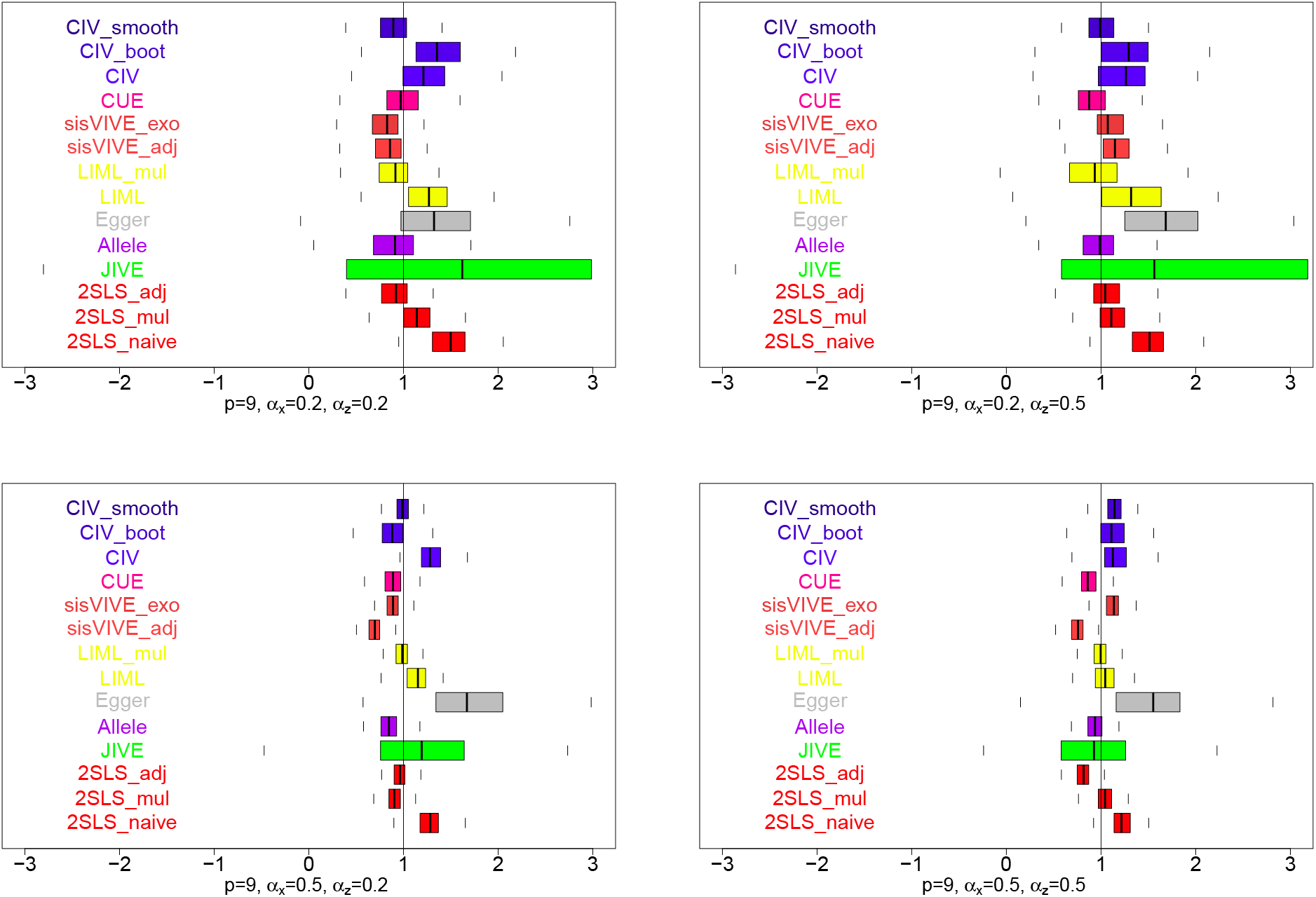
Boxplots of estimates of the causal effect estimates, *β*, from a one-sample set-up in simulation series III, with *p* = 9 instruments across 200 simulations, when true *β* = 1. The panels display results for different values of *α_x_* and *α_z_*.

**Figure 20:**
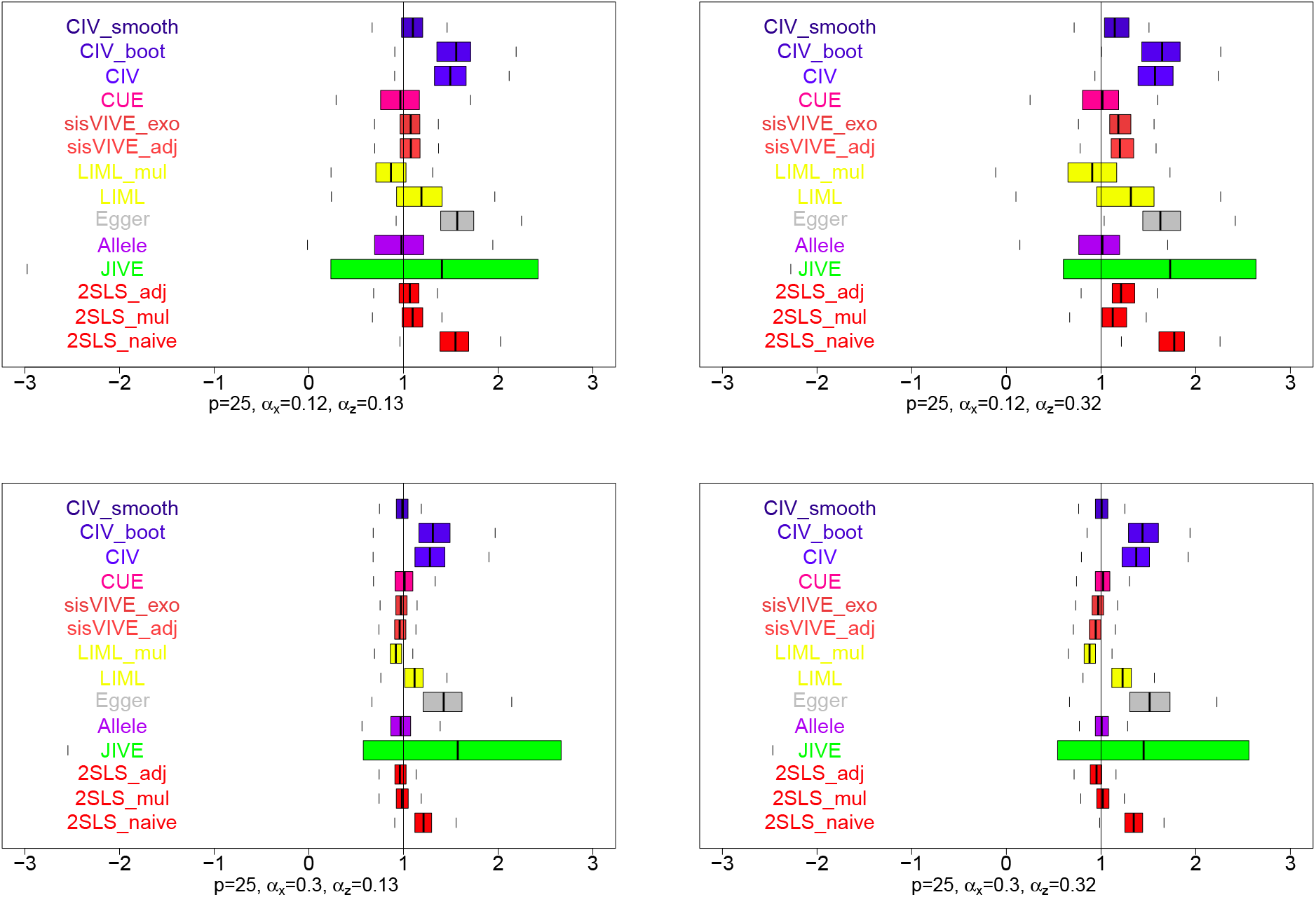
Boxplots of estimates of the causal effect estimates, *β*, from a one-sample set-up in simulation series III, with *p* = 25 instruments across 200 simulations, when true *β* = 1. The panels display results for different values of *α_x_* and *α_z_*

**Figure 21:**
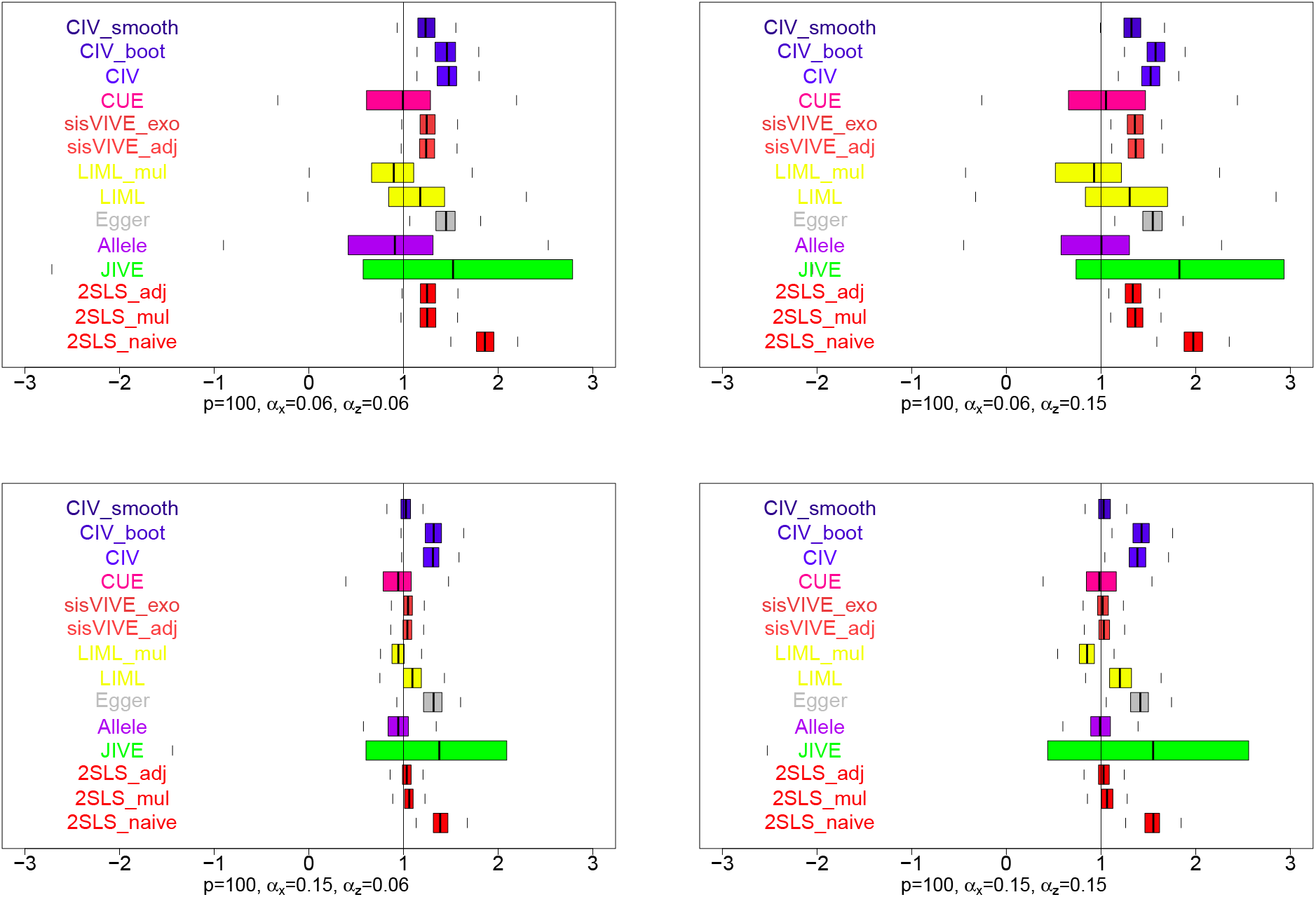
Boxplots of estimates of the causal effect estimates, *β*, from a one-sample set-up in simulation series III, with *p* = 100 instruments across 200 simulations, when true *β* = 1. The panels display results for different values of *α_x_* and *α_z_*

In our simulations, MR analyses were mostly restricted to the case of a single risk factor **X**, although most of the methods mentioned above can be extended in some way to allow for a multivariate **X**. *CIV* and *CIV_smooth* methods can be adapted to allow for multivariate **X** as seen in Equation 5, and the corresponding multiple solutions c can be used with multivariate 2SLS to infer the causal effect of **X** on **Y**. *sisVIVE* is also flexible to the dimension of **X** as seen in Equation (4). Generalized moment methods (e.g. *LIML, CUE*) and 2SLS can also be adapted to handle multiple risk factors together. However, the nature of *Egger* regression restricts this approach primarily to univariate risk factor analysis; users can only analyze multiple risk factors one-by-one.

In simulations, we conducted two-sample analyses, by separating the weight construction process and causal inference process onto two different samples, to evaluate the properties of applicable instrumental variable methods. In such two-sample approaches, winner’s curse, which may result in over-estimates of the predictive power of constructed instruments in one-sample analyses, is less likely to occur.

In the ADNI data analysis, we applied a two-stage approach in a similar way, by constructing instruments using only the control samples. These instruments were then used in the whole dataset to estimate causal effect of each individual biomarker while treating others as secondary phenotypes. However, we do realize this two-stage ADNI analysis is still not an ideal solution due to the retrospective nature of the sampling in this case-control study sampling. There are additional techniques, including inverse weighting with sampling probability (Monsees et al., 2009) and maximum likelihood (Lin and Zeng, 2009), that can be considered in the future to correct for the case-control sampling. It is also worth noting that we transformed the causal effect estimation problem with multiple phenotypes into multiple causal effect estimations, each with an individual phenotype – versus the rest – for comparison purposes, since only *CIV* can process multiple causal effect estimations simultaneously while both *Allele* and *sisVIVE* are designed for individual phenotypes. This “one versus the rest” strategy leaves room for improvement in future work. The result of ADNI analysis in this paper simply serve as a demonstration of our *CIV* methods, rather than a strong causal statement of Alzheimer’s disease.

In conclusion, this paper proposed new approaches (*CIV* and *CIV_smooth* methods) for conducting Mendelian randomization analyses when the pleiotropy is observed. Under the assumption of linear structural equation models, these approaches can be used to implement valid instruments selection and adjust causal effect estimation when potential pleiotropic phenotypes are measured. Our methods *(CIV* and *CIV_smooth*) perform consistently in simulations and real data analysis. Hopefully, these methods will help to make the MR analyses a much more common practice even when pleiotropy is observed.

### A Solution to the Constrained Instrumental Variable Problem

Let *M* = *X*(*X*^⊤^*X*)^−1^*X*^⊤^ then *c*^⊤^*G*^⊤^*Xv* = *c*^⊤^*G*^⊤^*X*(*X*^⊤^*X*)^−1^*X*^⊤^*Xv* = *c*^⊤^*G*^⊤^*MXv*,

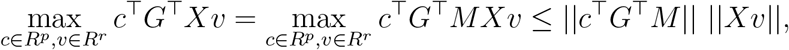

where 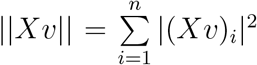 denotes the norm of vector *Xv* in the inner product space *R^p^*. The equality holds if and only if *Xv* and *MGc* are collinear. Let 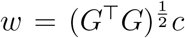 then the problem is equivalent to

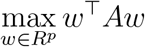

subject to conditions:

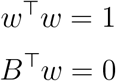

where 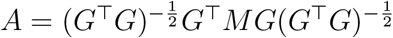 and 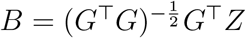.

If we have rank(*A*) = *p* ≥ rank(*Z*) = *k* (columns are uncorrelated), then there exists solution for *w* since this is a quadratic optimization problem with quadratic/linear constraints (Golub, 1973).

Consider the QR decomposition of *B*:

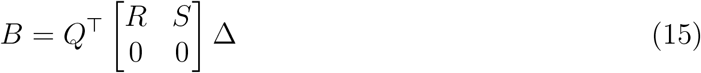

where *R* is a *k* by *k* upper triangular matrix with positive diagonal elements (thus invertible). *Q* is an orthogonal matrix. *S* is a *k* by *p* – *k* matrix and Δ represents the column permutation matrix (Gu and Eisenstat, 1996) to ensure that the diagonal elements of *R* are positive and non-increasing, i.e. *R* is invertible. *R* is then unique under these conditions (Golub and Van Loan, 1996).

Now we let 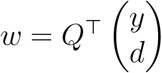 where *y* ∈ *R^k^*, *d* ∈ *R*^*p−k*^ and 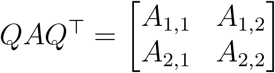.

The problem then becomes:

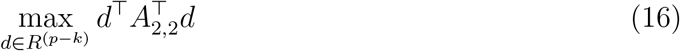

subject to conditions:

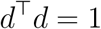

We now know that the solution for *d* is the eigenvector corresponding to the largest eigenvalue of *A*_2,2_. There are at most *p – k* eigenvectors.

In conclusion, when *n* > *p* the (unique) solution of the constrained instrumental variable (*CIV*) is 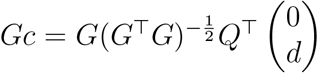, where *Q* is an orthogonal matrix defined by (15) and *d* is an eigenvector defined by (16).

### B Simulation III: Non-zero Null Hypothesis

